# Rapid aging and disassembly of actin filaments from two evolutionary distant yeasts

**DOI:** 10.1101/2025.11.21.689671

**Authors:** Ingrid Billault-Chaumartin, Hugo Wioland, Audrey Guillotin, Alphée Michelot, Antoine Jégou, Guillaume Romet-Lemonne

## Abstract

Actin is an essential cytoskeletal protein that is extremely well conserved across the tree of life. Similarities and differences in the way actin from different species self-assembles into filaments inform our understanding of its evolution. However, this basic knowledge is largely incomplete. Here, we address this issue by systematically characterizing assembly kinetics for actin from two yeast species that are five hundred million years apart in evolution, *Saccharomyces cerevisiae* and *Schizosaccharomyces pombe*, and compare them to the well-studied actin from rabbit skeletal muscle from which they diverged a billion years ago. By monitoring individual actin filaments under controlled conditions, we accurately quantified their barbed end elongation from ATP-actin monomers, their disassembly in the ADP·Pi and ADP states, as well as the release of inorganic phosphate, which marks their aging. We find that, in the ATP state, both yeast actins behave strikingly like mammalian actin at filament barbed ends. In contrast, yeast actin filaments in both the ADP·Pi and the ADP states depolymerize several-fold faster than their mammalian counterparts, and they release inorganic phosphate over 20-fold faster. We show that the absence of methylation on histidine 73 largely accounts for this faster aging of yeast actin filaments. We also reveal differences between the actins of the two yeasts. In particular, ADP-actin filaments depolymerize faster and are mechanically more flexible in *Saccharomyces cerevisiae* compared to *Schizosaccharomyces pombe*. Our findings suggest that actins expressed across species possess more diverse and specialized biochemical characteristics than previously recognized.

## Introduction

In living cells, actin filaments and actin-binding proteins (ABPs) form dynamic networks that support fundamental functions such as division, motility and polarity (Lappalainen et al., 2022; Pollard & Cooper, 2009). Filamentous actin (F-actin) results from the assembly of the monomeric, globular protein actin (G-actin) into polar filaments, with so-called barbed and pointed ends, the former being more dynamic. G-actin contains an ATP molecule which upon incorporation onto the filament quickly gets hydrolysed into ADP·Pi (Korn et al., 1987) The inorganic phosphate (Pi) is subsequently released, giving rise to ADP-actin. The 3 different nucleotide states of F-actin each have different assembly dynamics, both at the barbed and pointed end (Fujiwara et al., 2007; Pollard, 1986).

Actin is one of the most conserved proteins, ubiquitously expressed throughout the eukaryotic kingdom, and with homologs in Archaea (Gunning et al., 2015; Rodrigues-Oliveira et al., 2023). Some organisms, such as commonly studied yeasts, express only one actin gene. Others express multiple actin isoforms, such as mammals which typically have 6 different isoforms (Perrin & Ervasti, 2010) or plants which can express up to 21 distinct isoforms (Šlajcherová et al., 2012). The alpha-skeletal muscle actin isoform of rabbit (*Oc*ACTA1), or the 100% amino acid sequence-identical homolog from chicken, has become a standard in the field due to its high abundance in muscle and good purification yield (Spudich & Watt, 1971). Most of our knowledge on actin dynamics stems from *in vitro* studies using actin from this single gene.

In spite of actin being highly conserved, extrapolating results from *Oc*ACTA1 to other actin orthologs must be done with caution. Notably, heterologous actins are not systematically interchangeable (Boiero Sanders et al., 2022). For instance, substituting actin in *Saccharomyces cerevisiae* (*Sc*Act1) with other eukaryotic actins is lethal or leads to various defects *in vivo*. The severity of the phenotype correlates negatively with the degree of homology between the actins (Boiero Sanders et al., 2022). In particular, despite a 90% amino acid identity, the replacement of *Sc*Act1 with the *Schizosaccharomyces pombe* actin gene (*Sp*Act1) is not viable. Furthermore, within one given organism, different isoforms can support distinct functions and can exhibit distinct localizations (Boiero Sanders et al., 2020; Perrin & Ervasti, 2010; Šlajcherová et al., 2012).

The non-interchangeability of actins is certainly due in part to some ABPs being isoform-specific. For example, cofilin from *Leishmania major* does not bind to *Oc*ACTA1 filaments, in spite of Leishmania cofilin-actin filaments being strikingly similar to their mammalian counterparts (Kotila et al., 2022). Intrinsic differences in the assembly properties of actin isoforms likely contribute to their non-interchangeability as well. Such differences appear clearly as diverse actin isoforms are being studied. For example, actin filaments from Apicomplexan parasites, which are distant in evolution from mammals, turn over much faster due to faster disassembly rates and a higher critical concentration for assembly (Lu et al., 2019; Hvorecny et al., 2024). These intrinsic kinetic differences are key for our understanding of the actin cytoskeleton and its evolution, which is still very incomplete.

In that respect, the two distant yeasts *Saccharomyces cerevisiae* (*S. cerevisiae*) and *Schizosaccharomyces pombe* (*S. pombe*) are particularly interesting. They diverged from each other about half a billion years ago (Shen et al., 2020) and twice that long ago from metazoans (Dohrmann & Wörheide, 2017), thus probing an intermediate region in the eukaryotic tree. In addition, both have been used as model organisms for decades, are amenable to quick genetic manipulation and have well-characterized actin networks (Mishra et al., 2014; Kovar et al., 2011). While their actin assembly kinetics are known to differ from the standard *Oc*ACTA1, quantifications of reaction rates and systematic comparisons are lacking. For example, *Sc*Act1 and *Sp*Act1 filaments have been reported to assemble faster, but this feature is attributed to a more efficient filament nucleation, and elongation rates have not been measured (Buzan & Frieden, 1996; Ti & Pollard, 2011). Likewise, *Sp*Act1 and *Sc*Act1 filaments have been reported to release Pi rapidly, but no reaction rate has been measured (Ti & Pollard, 2011; Yao & Rubenstein, 2001).

Here, we aim to fill this gap, taking advantage of recent techniques for the production of actin (Hatano et al., 2018, 2020) and for the monitoring of their assembly dynamics at the single filament level (Jégou et al., 2011; Wioland et al., 2022) using a low fraction of fluorescently labeled nucleotides (Colombo et al., 2021). We find that, in spite of their evolutionary distance, the two yeasts share highly similar ATP-actin dynamics to mammalian skeletal muscle actin, but accelerated ADP·Pi and ADP-actin barbed end depolymerization rates, in addition to a strikingly accelerated Pi-release rate. The accelerated Pi-release rate is largely explained by the absence of histidine 73 methylation, which is ubiquitous in mammalian actins but absent in yeast actins.

## Results

### ATP-actin from *Saccharomyces cerevisiae* and *Schizosaccharomyces pombe* polymerize at the same rate as mammalian actin

*Saccharomyces cerevisiae* (*S. cerevisiae*) and *Schizosaccharomyces pombe* (*S. pombe*) have single actin genes (coding for the proteins *Sc*Act1 and *Sp*Act1, respectively) that share close amino acid sequence identity to the 6 mammalian actin genes, reflecting the exceptional conservation of the actin gene in general (Figure S1). In particular, they respectively share 87 and 89% amino-acid sequence identity with the α-skeletal muscle *Oryctolagus cuniculus* (rabbit) homolog, *Oc*ACTA1, used in most *in vitro* actin characterization studies (Figure 1A and Table 1). *Sc*Act1 and *Sp*Act1 are only slightly more similar to each other than they are similar to *Oc*ACTA1, with 90% amino-acid sequence identity. In spite of these strong similarities in sequence, substituting one ortholog for another is lethal *in vivo* (Boiero Sanders et al., 2022).

**Figure 1.**
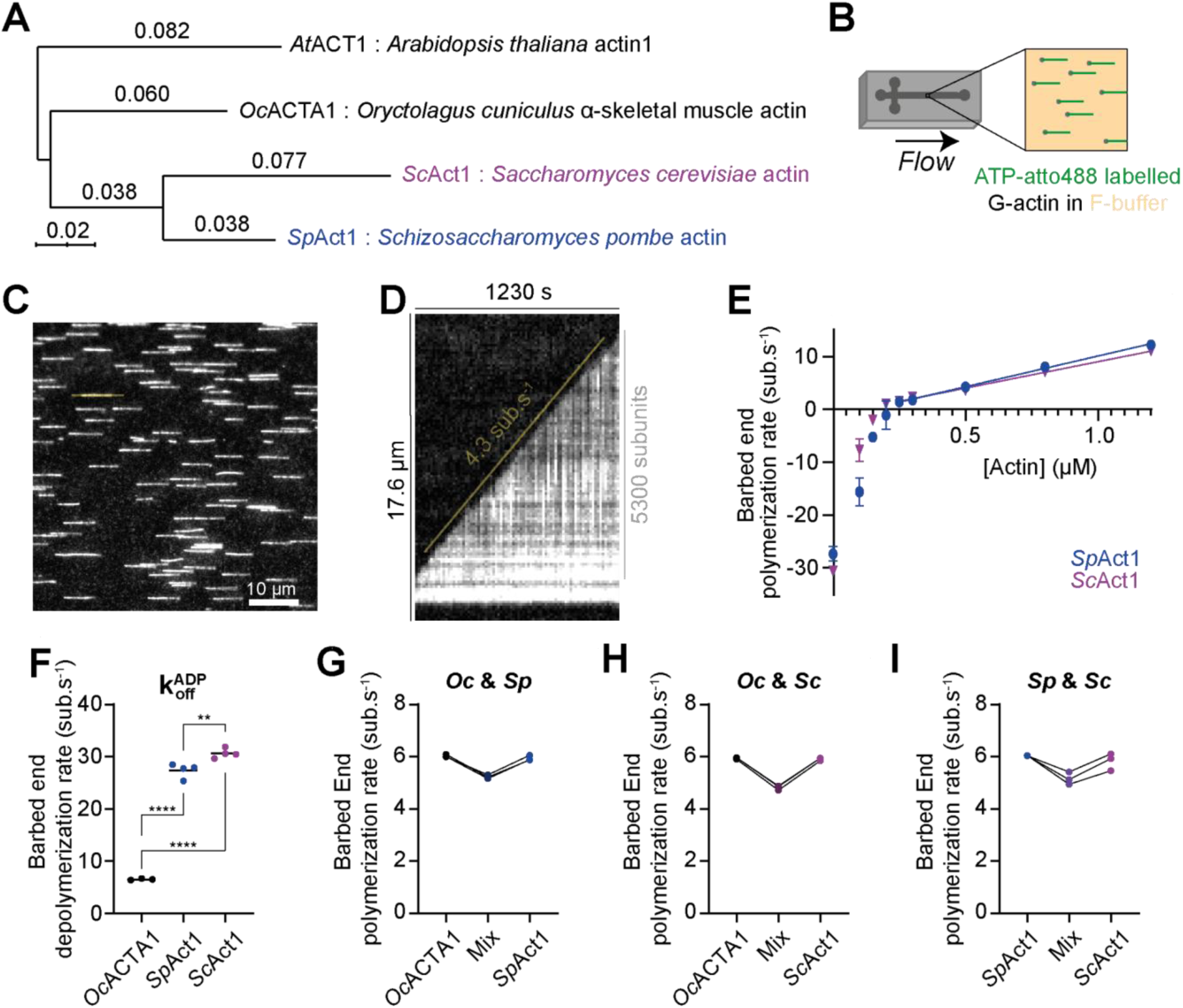
*Saccharomyces cerevisiae* and *Schizosaccharomyces pombe* actins exhibit indistinguishable ATP-actin dynamics from rabbit skeletal muscle actin but faster ADP-actin depolymerization rates. **A.** Phylogenetic tree of the 3 actins used in this study, *Saccharomyces cerevisiae* actin *Sc*Act1 (Uniprot P60010), *Schizosaccharomyces pombe* actin *Sp*Act1 (Uniprot P10989) and *Oryctolagus cuniculu*s (rabbit) alpha skeletal muscle actin *Oc*ACTA1 (Uniprot P68135). *Arabidopsis thaliana* actin 1 *At*ACT1 (Uniprot P0CJ46) was used as an external group. Scale bar unit is the average number of substitutions per amino-acid. **B.** Method used to study the dynamics of actin filaments in TIRF-assisted microfluidics experiments. ATP-ATTO488 G-actin (green) is flowed into a PDMS microfluidic chamber and polymerized in F-buffer (yellow) onto spectrin-actin seeds (grey) adsorbed onto the surface. This results in filaments anchored to the surface by their pointed end and whose dynamic barbed end can be monitored using TIRF microscopy. **C.** Example of a cropped field of view obtained with the method sketched in B using 0.5 µM *Sp*Act1 and 0.25 µM ATP-ATTO488. Movie S1 shows this cropped field of view over the course of the experiment. **D.** Kymograph of the filament outlined in C (yellow line) over the course of the acquisition. The elongation rate is obtained from the angle formed by the line following the filament barbed end over time (yellow line) and the abscissa. **E.** Plot of the barbed end polymerization rate as a function of the actin concentration for both *Sp*Act1 (blue) and *Sc*Act1 (magenta). Each point is the mean of at least 4 independent experiments where the elongation rate of 30 filaments was quantified as in D, and error bars are standard deviations. A linear regression for points from 0.25 µM to 1.2 µM actin was used to extract k_on_ ^ATP^ as the slope and k_off_^ATP^ as the y-intercept (Table 3). **F.** The rates k_off_^ADP^ correspond to the values at 0 µM actin in E for *Sc*Act1 and *Sp*Act1 and 0 mM phosphate in Figure 2C for *Oc*ACTA1 (Table 3), measured after polymerizing the filaments. Each point is the mean of an independent replicate, and black bars are means of the replicates for a given condition. Indicated significances are from unpaired student t-tests. (**: p≤10^-2^ ; ****: p≤10^-4^). **G-I.** Plots of the barbed end polymerization rate for pure solutions of *Oc*ACTA1 (black – G and H), *Sp*Act1 (blue – G and I), *Sc*Act1 (magenta – H and I) and their 50/50 respective mixes. Please note that the pure and mixed solutions are polymerized simultaneously in these experiments (see Figure S2A) to ensure identical experimental conditions. Each point is the mean of an independent replicate where the polymerization rate of 30 filaments was quantified, and points across categories stemming from the same acquisition are linked.

**Table 1.** Pairwise sequence identity of the actin sequences considered in. **Figure 1A**

|  | *A. thaliana* | *O. cunigulus* | *S. pombe* | *S. cerevisiae* |
| --- | --- | --- | --- | --- |
| *A. thaliana* | 100.0 | 87.80 | 87.47 | 83.47 |
| *O. cunigulus* | 87.80 | 100.0 | 88.80 | 86.67 |
| *S. pombe* | 83.47 | 88.80 | 100.0 | 89.60 |
| *S. cerevisiae* | 83.47 | 86.67 | 89.60 | 100.0 |

To study the actin polymerization properties of the two divergent yeast actins of *S. cerevisiae* and *S. pombe*, we took advantage of a recently developed system that allows for the expression of recombinant eukaryotic actins in *Pichia pastoris* (Hatano et al., 2018, 2020). We purified both actins using this system in the presence of the N-terminal acetylation and in the absence of histidine 73 methylation, as is the case *in vivo* (Table 2). We monitored actin filaments using a low fraction of ATP-ATTO488 (Colombo et al., 2021), a method that is useful when the direct chemical labeling of actin is challenging, and that does not alter the rates at which filaments elongate (Colombo et al., 2021) and release Pi (Schahl et al., 2025). We then studied their barbed end polymerization kinetics by microfluidics-assisted TIRF microscopy (Figure 1B; Jégou et al., 2011; Wioland et al., 2022) at different actin concentrations. Experiments were carried out at 25°C, pH 7.4 (Methods). With this method, spectrin-actin seeds are anchored to the surface and the filaments grown from them are conveniently aligned by the flow (Figure 1C), easing the monitoring of their barbed end elongation over time (Figure 1D, Movie S1). This allowed us to measure the elongation rate as a function of the concentration of actin monomers (Figure 1E).

**Table 2.**
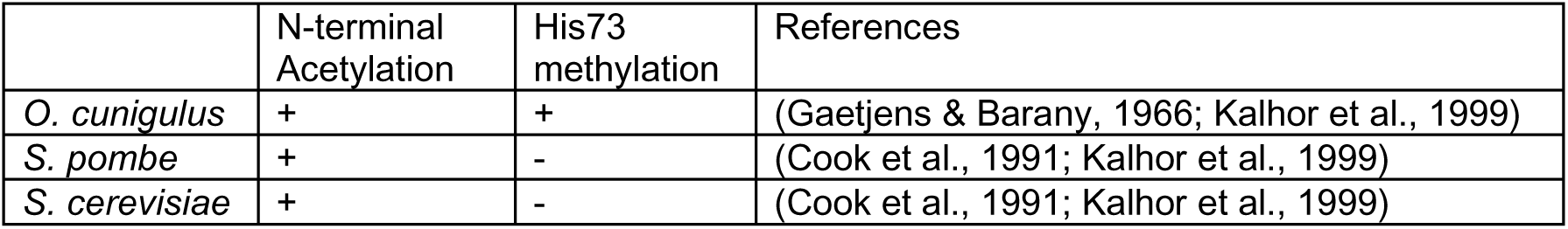
*In vivo* presence of the two major actin PTMs in relevant actins.

At high actin concentrations, polymerization is fast enough to consider that the barbed end is mostly in the ATP state, and a linear fit yields the on- and off-rate constants (k_on_^ATP^ and k_off_^ATP^) of ATP-actin at the barbed end (Figure 1E). Strikingly, these reaction rates for *Sp*Act1 and *Sc*Act1 are very similar to those published for *Oc*ACTA1 (Figure 1E, Table 3; Pollard, 1986; Jégou et al., 2011) with k_on_ = 11.6±0.7, 10.0±0.7 and 11.6±1.2 µM^-1^ s^-1^, respectively, and k_off_ = 1.5±0.5, 1.0±0.5 and 1.4±0.8 s^-1^, respectively. To our knowledge, these rate constants have not been previously measured, but our results are consistent with earlier studies in bulk assays on *S. cerevisiae* where faster polymerization was observed but attributed to nucleation rather than elongation (Buzan & Frieden, 1996), and bulk assays on *S. pombe* actin where elongation rates were not quantified but proposed to be similar to that of mammalian actin (Takaine & Mabuchi, 2007; Ti & Pollard, 2011).

**Table 3.**
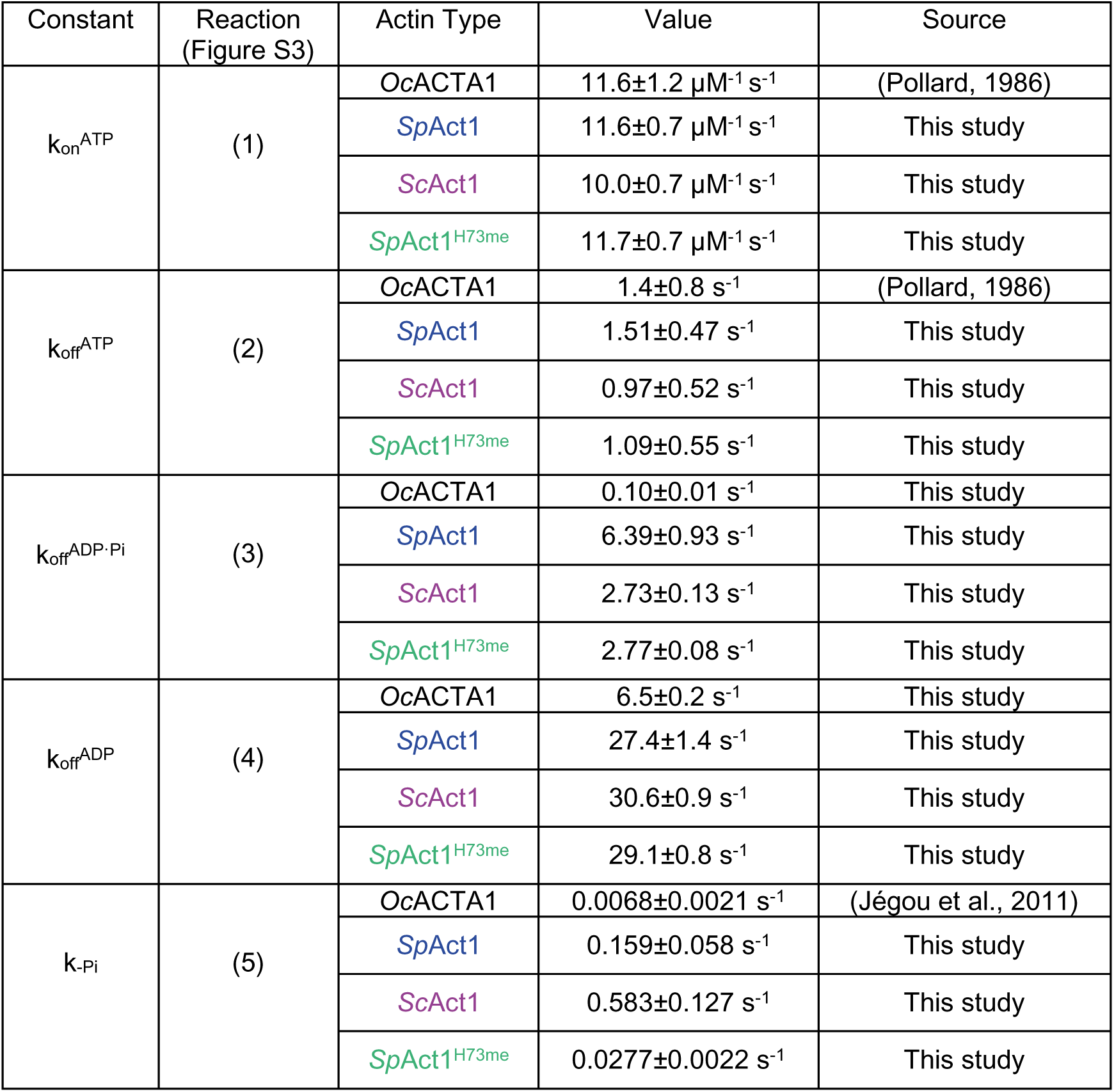
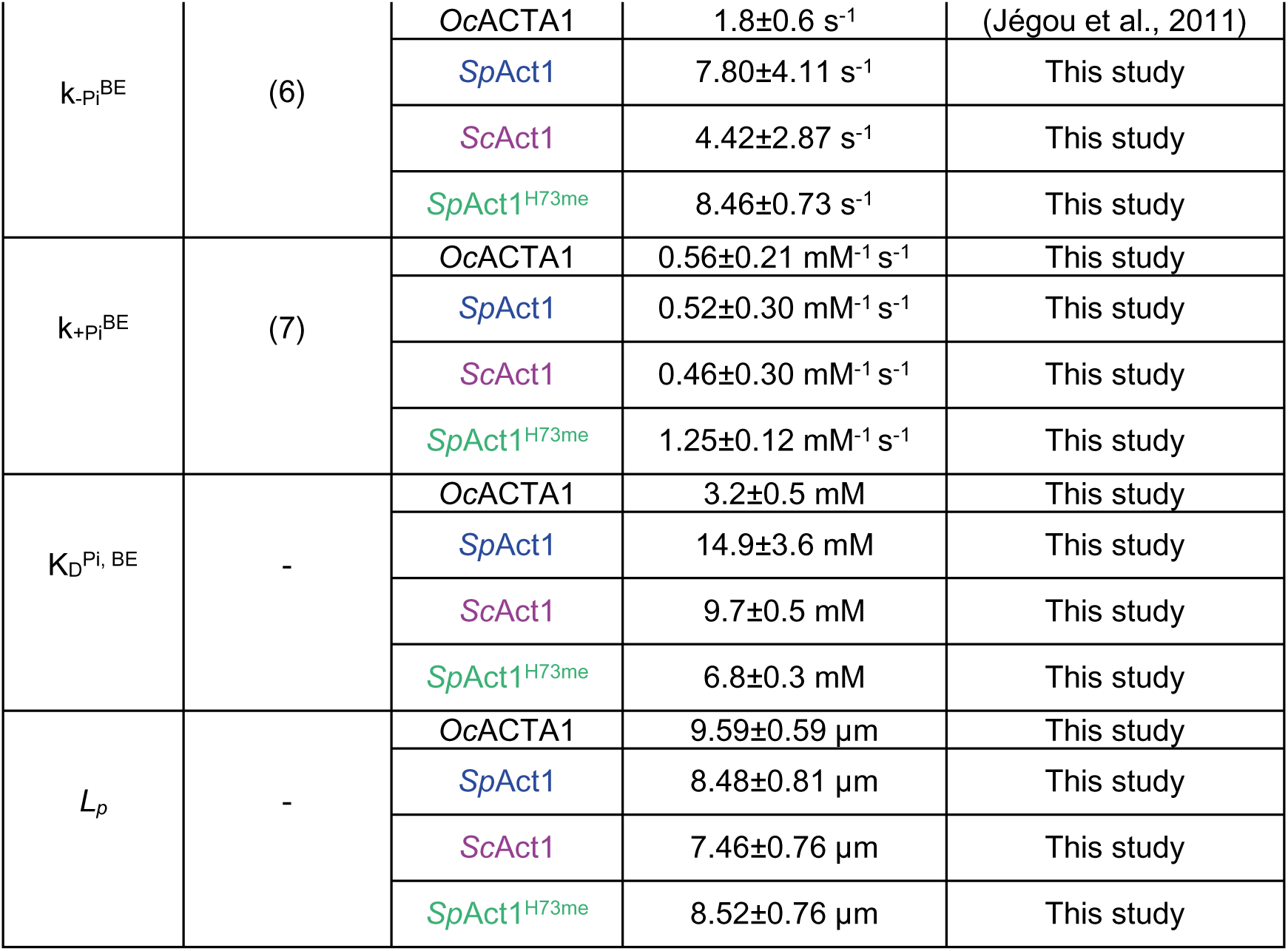
Results Summary For all reaction rates measured in this study, the indicated values are means ± standard deviations (over N≥3 repeats), except (1) and (2) which result from a linear fit and indicate the 95% confidence interval. See Figure S3 for a scheme showing the different reaction rates.

Taken together, these results show that the two divergent yeasts *S. cerevisiae* and *S. pombe* share nearly identical ATP-actin barbed end dynamics to each other and to the well-studied metazoan actins. This was far from expected, since a single point mutation on an actin isoform is enough to greatly alter its elongation rate (Greve et al., 2024). In spite of their nearly identical rate constants, we found that these actin orthologs were not fully interchangeable: mixing equal volumes of *Oc*ACTA1/*Sp*Act1, *Oc*ACTA1/*Sc*Act1, or *Sp*Act1/*Sc*Act1 solutions resulted in filaments elongating 13-19% slower than from the unmixed orthologs (Figures 1G-I and S2A-B). However, filaments made with actin from two different species did not appear to be substantially more fragile (Figure S10).

### Actin filaments from Saccharomyces cerevisiae and Schizosaccharomyces pombe disassemble much faster than mammalian actin filaments

At low actin concentrations, we found that *Sp*Act1 and *Sc*Act1 filaments depolymerized rapidly and at similar rates (Figure 1E). In this regime, the addition of subunits to the barbed end is less frequent than the rate of ATP hydrolysis in F-actin (0.3 s^-1^ for rabbit skeletal muscle actin; Blanchoin & Pollard, 2002) and the terminal subunits can be in different nucleotide states. In the absence of G-actin in solution, filaments reach a steady-state where they depolymerize as ADP-actin, at a rate k_off_^ADP^. We found that the two yeast actins used in this study depolymerized roughly 4.5-fold faster than rabbit alpha-skeletal actin (Figure 1E-F, Table 3): k_off_^ADP^ = 27.4±1.4 and 30.6±0.9 s^-1^ for *Sp*Act1 and *Sc*Act1, respectively, compared to 6.5±0.2 s^-1^ for *Oc*ACTA1 in the same conditions (similar to published values for *Oc*ACTA1 (Fujiwara et al., 2007; Wioland et al., 2019)).

By comparing recombinant and native actins for both *S. cerevisiae* and mammals, we verified that the differences in depolymerization did not come from potential post-translational modifications (Figure S5). We also verified that increasing the temperature did not affect the ranking of the different actins in terms of depolymerization rate (Figure S6).

We next sought to investigate the depolymerization rate of *S. cerevisiae* and *S. pombe* ADP·Pi-actin. The ADP·Pi-actin situation is more complex because the barbed end subunit is known, at least in mammalian actin, to release its inorganic phosphate (Pi) far more rapidly than subunits within the filament (Jégou et al., 2011; Oosterheert et al., 2023). As a consequence, an ADP·Pi-actin filament depolymerizes at a rate v_ADP·Pi_ which combines two routes, as subunits can either dissociate as ADP·Pi-actin at a rate k_off_^ADP·Pi^, or release their Pi and then dissociate as ADP-actin at a rate k_off_^ADP^ (Fig S3). In order to determine k_off_^ADP·Pi^, one can force dissociation to follow the first route. To do so, we polymerized filaments in phosphate buffer, then switched to the same phosphate buffer condition without actin (Figure 2A-B, Movie S2). The higher the concentration of Pi, the closer the barbed end subunits are to being fully in the ADP·Pi state, and the depolymerization rate eventually plateaus at k_off_^ADP·Pi^ when all subunits dissociate as ADP·Pi-actin (Figure 2C). We used the depolymerization rate measured at our highest Pi concentration as an estimate of this plateau value (Figure 2D, Table 3), yielding k_off_^ADP·Pi^ = 6.4±0.9 and 2.7±0.1 s^-1^ for *Sp*Act1 and *Sc*Act1, respectively, and 0.10±0.01 s^-1^ for *Oc*ACTA1 (similar to the published value (Fujiwara et al., 2007)). Unlike the disassembly of ADP-actin subunits, the disassembly rates of ADP·Pi-*Sp*Act1 and *Sc*Act1 differ substantially. They are also much larger than for *Oc*ACTA1: about 64-fold faster for *Sp*Act1 and 27-fold faster for *Sc*Act1.

**Figure 2.**
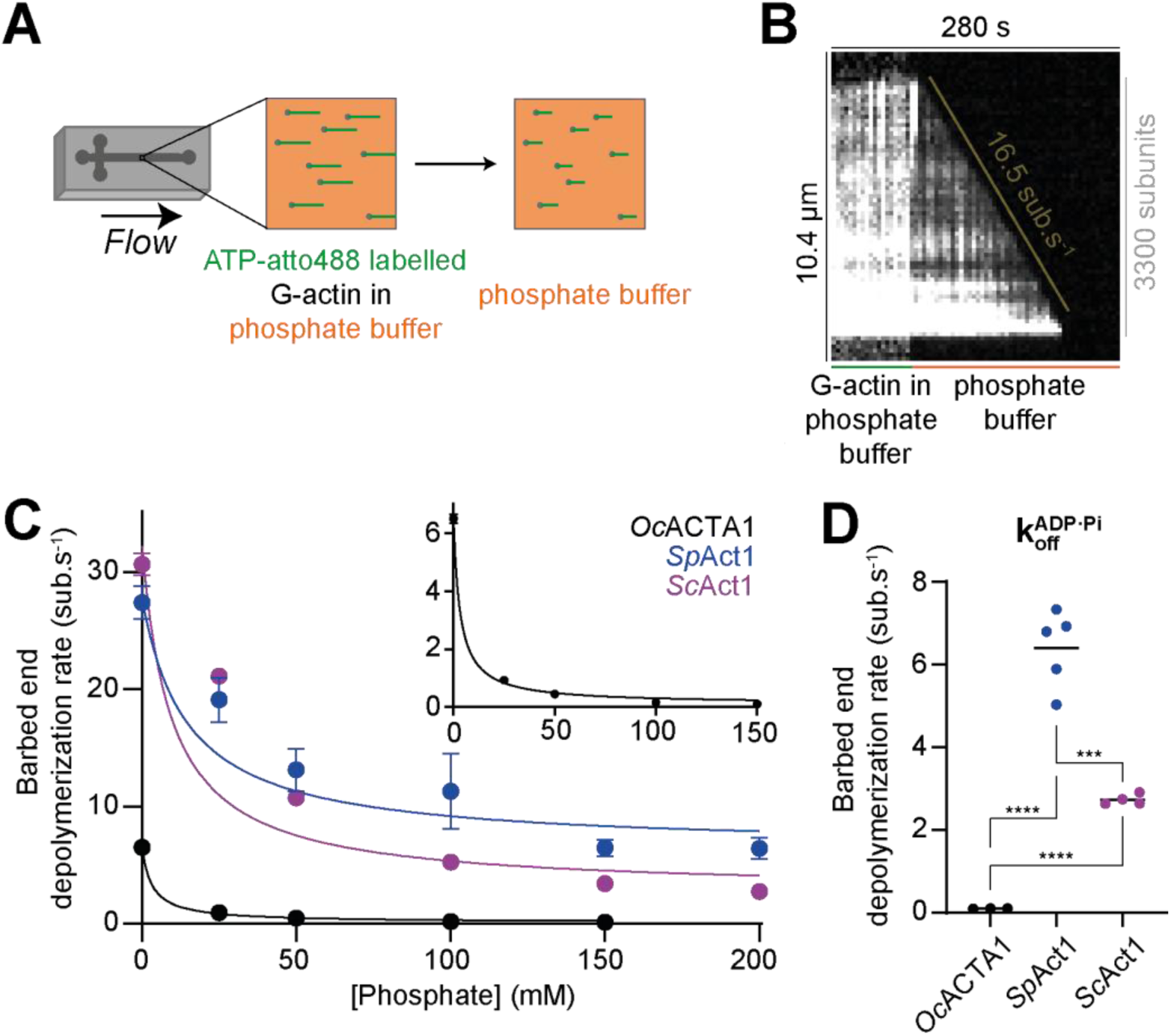
*Saccharomyces cerevisiae* and *Schizosaccharomyces pombe* ADP·Pi-actin filaments depolymerize fast. **A.** Method used to study the depolymerization of ADP·Pi-actin filaments in TIRF-assisted microfluidics experiments. ATP-ATTO488 G-actin (green) is flowed into a PDMS microfluidic chamber and polymerized in phosphate-buffer (orange) onto spectrin actin seeds (grey). The filaments are then exposed to the same phosphate buffer without actin and their depolymerization is monitored. **B.** Kymograph of an actin filament over the course of an experiment as in A using 1 µM *Sp*Act1, 0.5 µM ATP-atto488 and 25 mM phosphate (see Movie S2). **C.** Plot of the barbed end depolymerization rate as a function of the phosphate concentration for *Sp*Act1 (blue), *Sc*Act1 (Magenta) and *Oc*ACTA1 (black). Inset: rescaled plot of the *Oc*ACTA1 data. Each point is the mean of at least 3 independent experiments where the elongation rate of at least 15 filaments was quantified as in B and error bars are standard deviations. Solid lines are fits by hyperbolic functions (see Methods). **D.** Depolymerization rates k_off_^ADP·Pi^ correspond to measurements at 200mM phosphate for *Sp*Act1 and *Sc*Act1, and 150 mM phosphate for *Oc*ACTA1, in C (Table 3). Each point is the mean of an independent replicate, and black bars are means of the replicates for a given condition. Indicated significance is from an unpaired student t-test (***: p≤10^-3^; ****: p≤10^-4^)

When filaments depolymerize in phosphate buffer, Pi rapidly binds and unbinds the nucleotide pocket of the terminal subunit at the barbed end, and the titration curves of Figure 2C can be used to determine the corresponding equilibrium dissociation constant, K_D_^Pi, BE^ (Methods). We find that Pi has a weaker affinity for the barbed ends of yeast actin filaments, compared to *Oc*ACT1 (Table 3).

Changing the phosphate concentration has an effect on the ionic strength of the solution and on its ability to solubilize proteins (ion-specific, Hofmeister effects), which could themselves potentially affect the depolymerization rate. To control for these possible effects, we measured *Sp*Act1 and *Sc*Act1 depolymerization rates with increasing concentrations of KCl and with increasing concentrations of Na_2_SO_4_ (SO_4_^2-^ being similarly kosmotropic as PO_4_^3-^, Mazzini & Craig, 2017). We found that they had a moderate impact on both *Sp*Act1 and *Sc*Act1 (Figure S4A-B).

Taken together, these results show that ADP- and ADP·Pi-actin filaments for both *S. cerevisiae* and *S. pombe* depolymerize strikingly faster than for mammalian skeletal muscle actin.

### Actin filaments from *Saccharomyces cerevisiae* and *Schizosaccharomyces pombe* release their inorganic phosphate much faster than mammalian actin filaments

An important aspect of actin dynamics is the transition from the ADP·Pi to the ADP state, which makes the filaments less stable and plays an important role in controlling the binding of regulatory proteins (Blanchoin & Pollard, 1999; Cai et al., 2007; Zimmermann et al., 2015). Hence, we next sought to investigate how fast yeast actins release their inorganic phosphate (Pi). For that, we polymerized filaments in phosphate buffer, as before, but then switched to a buffer without phosphate for depolymerization (Figure 3A). Because k_off_^ADP·Pi^ is lower than k_off_^ADP^, we observed an increase of the depolymerization rate over time, as Pi is progressively released from actin subunits of the filament (Figure 3B). By analyzing this acceleration (Figure S9) we can extract the Pi-release rate k_-Pi_ (Jégou et al., 2011; Kotila et al., 2022; Oosterheert et al., 2023; Methods).

**Figure 3.**
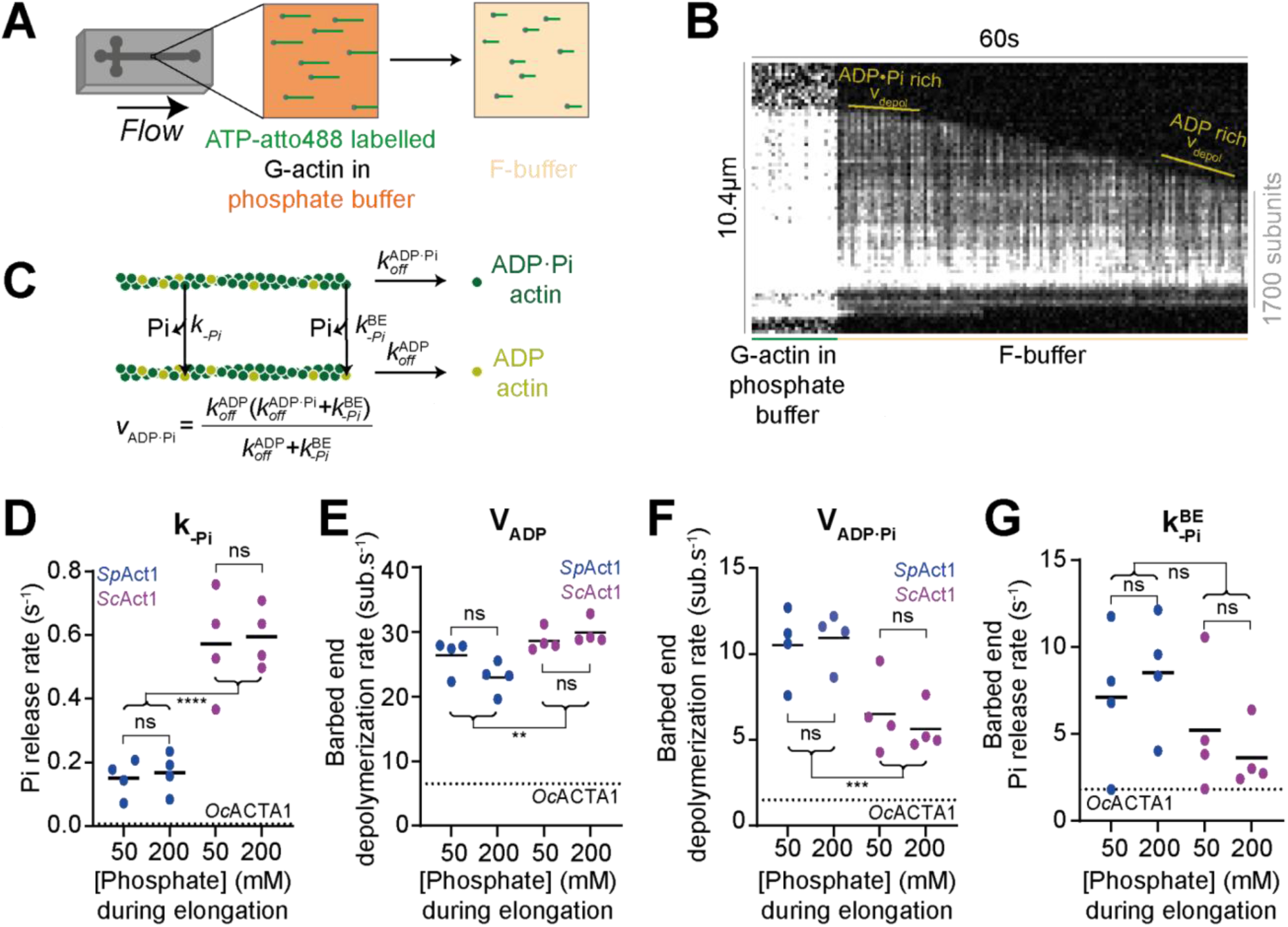
*Saccharomyces cerevisiae* and *Schizosaccharomyces pombe* actin filaments release inorganic phosphate faster than rabbit alpha-skeletal actin filaments. **A.** Method used to study the phosphate release rate of actin filaments in TIRF-assisted microfluidics experiments. ATP-ATTO488 G-actin (green) is flowed into a PDMS microfluidic chamber and polymerized in phosphate-buffer (orange) onto spectrin actin seeds (grey). The filaments are then exposed to F-buffer (yellow) without actin and depolymerization is monitored over time. **B.** Example of a kymograph of an actin filament over the course of an experiment as in A using 0.8 µM *Sp*Act1, 0.4 µM ATP-atto488 and 50 mM phosphate during elongation. Depolymerization takes place in F-buffer (no phosphate), at a rate that accelerates as the ADP·Pi content of the filament decreases. **C.** Scheme explaining the different rates measured in this experiment. The depolymerization rate V_ADP·Pi_ is in subunits per second. **D-G.** Plot of the phosphate release rate k_-Pi_ (D), the depolymerization rate V_ADP_ at the end of the experiment (E), the depolymerization rate V_ADP·Pi_ at the beginning of the experiment (F) and the barbed end phosphate release rate k_-Pi_^BE^ (G), for *Sp*Act1 (blue) and *Sc*Act1 (magenta) following a polymerization at 50 and 200 mM phosphate. Dotted lines are the values obtained for *Oc*ACTA1 in Jégou et al., 2011 (D, G) or measured in this study (E, F), for reference. Each point in (D, E, F) is the result of the fit of the depolymerization rate as a function of time, averaged over at least 40 filaments (Figure S9). Each point in (G) is calculated based on the values of V_ADP·Pi_ (F), k_off_^ADP·Pi^ (Figure 2D) and k ^ADP^ (Figure 1F) and using the equation in (C) which accounts for the two routes ADP·Pi-actin can follow to dissociate from the filament barbed end. Black bars are means of the replicates for a given condition. Indicated significances are from unpaired student t-tests (ns: p>0.05 ; **: p≤0.01 ; ***: p≤10^-3^ ; ****: p≤10^-4^).

We find that Pi release from F-actin is 23-fold faster for *S. pombe* and 86-fold faster for *S. cerevisiae*, compared to mammals: k_-Pi_ = 0.16±0.06 s^-1^ and 0.58±0.13 s^-1^ for *Sp*Act1 and *Sc*Act1, respectively (Figure 3D, Table 3), compared to 0.0068±0.0021 s^-1^ for *Oc*ACTA1 (rate from (Jégou et al., 2011), in agreement with (Melki et al., 1996; Fujiwara et al., 2007; Oosterheert et al., 2023)). Pi release in the two yeast actins differ, as it is 4-fold faster for *Sc*Act1 than *Sp*Act1. Our 23-fold increase for *Sp*Act1 compared to *Oc*ACTA1 is consistent with earlier bulk assays which estimated a 10- to 100-fold increase, lacking in precision due to a limited temporal resolution (Ti & Pollard, 2011).

Analyzing the acceleration of depolymerization over time (Figures 3B, S9, Methods) also yields the depolymerization rate of the ADP·Pi-actin filament, V_ADP·Pi_, corresponding to the beginning of the depolymerization, as well as the depolymerization rate of the ADP-actin filament, V_ADP_, corresponding to later times when the actin subunits have all released their Pi. By definition, V_ADP_ is identical to k_off_^ADP^, and the values we find (24.6±3.0 s^-1^ for SpAct1 and 29.3±1.8 s^-1^ for ScAct1, Figure 3E) match our previous measurements of k_off_^ADP^ (Figure 1F).

In contrast, V_ADP·Pi_ is faster than k_off_^ADP·Pi^, as revealed previously for rabbit skeletal muscle actin (Jégou et al., 2011). This is because, as mentioned earlier (Figure S3), actin subunits at the barbed end release their Pi at a rate k_-Pi_^BE^, faster than the rest of the filament. Thus, the depolymerization of ADP·Pi-actin can happen via two routes: (1) the ADP·Pi subunit dissociates from the barbed end as ADP·Pi-actin or (2) it first releases its Pi and then dissociates as ADP-actin (Figure 3C). We found that V_ADP·Pi_ is notably faster for *Sp*Act1 (V_ADP·Pi_ = 10.7±1.7 s^-1^) than *Sc*Act1 (V_ADP·Pi_ = 6.1±1.8 s^-1^, Figure 3F). Knowing V_ADP·PI_, k_off_^ADP·Pi^ and k_off_^ADP^, we can compute k_-Pi_^BE^ which is indeed much faster than k_-Pi_ (Figure 3G, Table 3). We find that k_-Pi_^BE^ is almost twice as fast for *Sp*Act1 (7.8±4.1 s^-1^) than for *Sc*Act1 (4.4±2.9 s^-1^), though with a weak statistical significance (p = 0.0554), and both are markedly faster than for *Oc*ACTA1 (1.8±0.6 s^-1^).

Since we have determined the equilibrium dissociation constant for Pi at the barbed end, K_D_^Pi, BE^ (Figure 2C), we can use it to compute the on-rate constant k_+Pi_^BE^=k_-Pi_^BE^/K_D_^Pi, BE^. We find very similar values for the three actin species (Table 3).

To verify that the actin filaments were indeed fully in the ADP·Pi state when switching to depolymerizing conditions, we repeated the experiment with different concentrations of phosphate buffer. Whether the filaments were polymerized in the presence of 50 or 200 mM phosphate, we found no significant difference in the rates we extracted from their subsequent depolymerization (Figure 3D-G). This result indicates that the bulk of the filament is already saturated with Pi at 50 mM (unlike the barbed end, Figure 2C), and validates our assay.

Together, these results show that *S. pombe* and *S. cerevisiae* actin filaments have faster Pi-release rates than mammalian actin filaments, both from subunits inside the filament and at the barbed end.

### The faster Pi-release rate of yeast actins is largely due to the absence of histidine 73 methylation

We next sought to determine why the Pi-release rates of *Sc*Act1 and *Sp*Act1 are faster than for *Oc*ACTA1. An appealing hypothesis is the known absence of the histidine 73 methylation in yeast, an essential and ubiquitous post-translational modification (PTM) in metazoan actins (Figure 4A, Table 2), whose removal from β rabbit actin was recently shown to accelerate Pi release (Schahl et al., 2025). In *S. cerevisiae*, mutations of histidine 73 were shown to affect actin assembly *in vitro* (Yao et al., 1999) and recent high-resolution cryoEM structure suggests that the hydrogen bonds in the vicinity of histidine 73 may affect Pi release (Stevenson et al., 2025).

**Figure 4.**
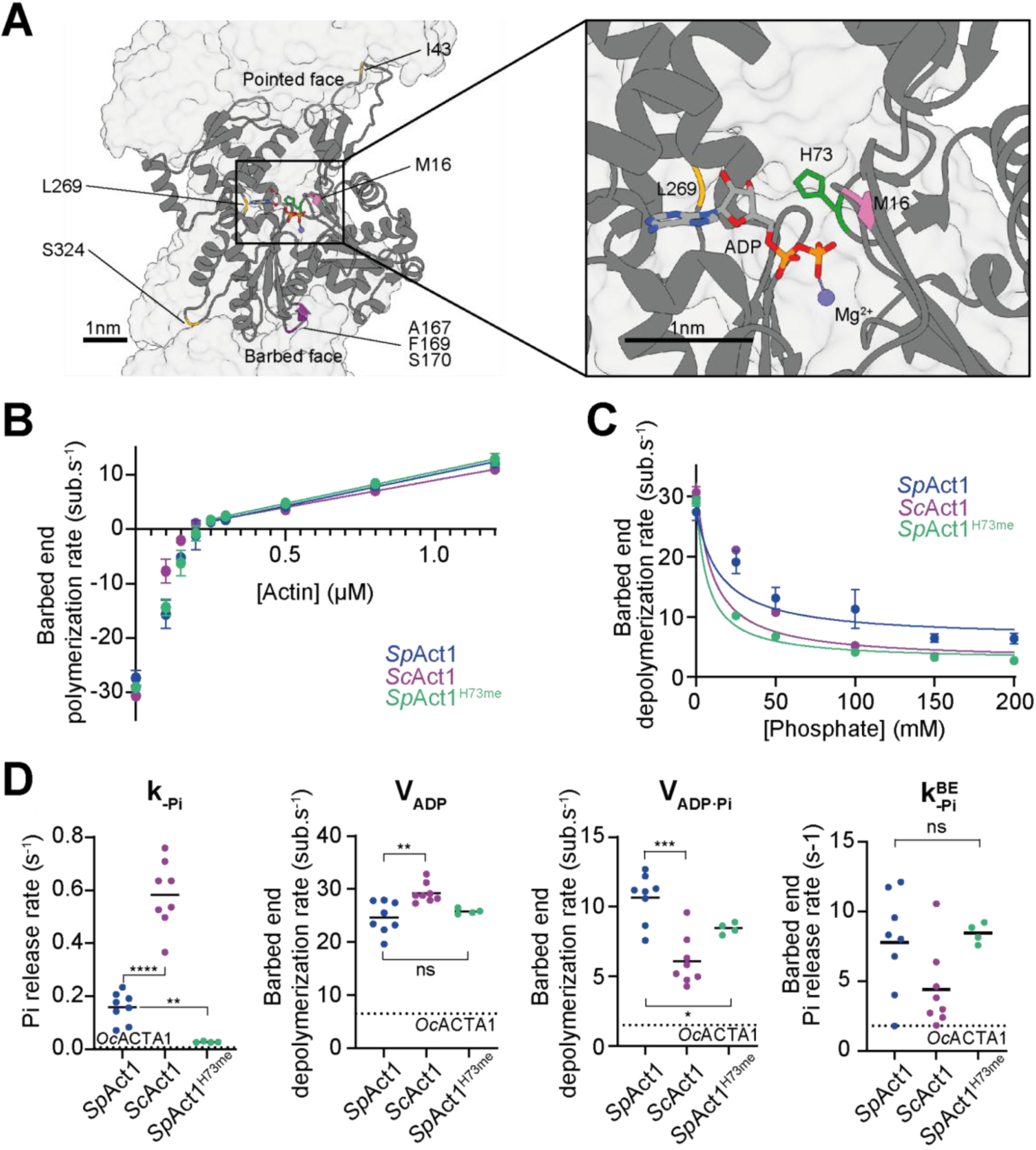
Histidine 73 methylation of *Schizosaccharomyces pombe* actin slows down Pi-release rate. **A.** Backbone of an *S. cerevisiae* F-actin subunit (PDB 9go5, from Stevenson et al., 2025) within the filament. The unmethylated histidine 73 (H73) residue is highlighted in green. Methionine 16 (M16), in pink, is present in *S. cerevisiae*, *S. pombe* and beta/gamma mammalian actins, but is replaced by leucine 16 in mammalian alpha-skeletal actin. In yellow are residues isoleucine 43 (I43), leucine 269 (L269) and serine 324 (S324), which are the same in *S. pombe* and *S. cerevisiae*, but are changed to valine 43, methionine 269 and threonine 324 in alpha-skeletal, beta and gamma cytoplasmic actins in mammals. Located in the W-loop, residues alanine 167 (A167), phenylalanine 169 (F169) and serine 170 (S170) in *S. cerevisiae* differ from their counterparts in both *O. cuniculus* and *S. pombe* and are in Magenta. **B.** Plot of the barbed end polymerization rate as a function of the actin concentration for *Sp*Act1^H73me^ (Green). Values for *Sp*Act1 (Blue) and *Sc*Act1 (Magenta) are a duplicate of Figure 1E and are added for reference. Each point is the mean of at least 4 independent experiments where the elongation rate of 30 filaments was quantified and error bars are standard deviation. A linear regression for points from 0.25 µM to 1.2 µM actin was used to extract k_on_^ATP^ as the slope and k_off_^ATP^ as the -Y interception (Table 3). The values at 0 µM actin give the k_off_^ADP^ (Table 3). **C.** Plot of the barbed end depolymerization rate as a function of the phosphate concentration for *Sp*Act1^H73me^ (Green). Values for *Sp*Act1 (Blue) and *Sc*Act1 (Magenta) are a duplicate of Figure 2C and are added for reference. Each point is the mean of at least 4 independent experiments where the elongation rate of at least 15 filaments was quantified, and error bars are standard deviation. Data at 200 µM phosphate was used for k_off_^ADP·Pi^ (Table 3). Solid lines are fits by hyperbolic functions, yielding the equilibrium dissociation constant K_D_ ^Pi, BE^ (see Methods). **D.** Plots of the phosphate release rate k_-Pi_, the depolymerization rate V_ADP_, the depolymerization rate V_ADP·Pi_ and the barbed end phosphate release rate k_-Pi_^BE^, for *Sp*Act1 (blue), *Sc*Act1 (magenta) and *Sp*Act1^H73me^(Green). Data for *Sp*Act1 and *Sc*Act1 are pooled data from Figure 3D-G included for comparison. Because polymerizing at 50 or 200 mM phosphate had no impact for *Sp*Act1 and *Sc*Act1, data for *Sp*Act1^H73me^ were extracted from experiments done at 50 mM exclusively. Dotted lines are the values obtained for *Oc*ACTA1 in (Jégou et al., 2011) (for k_-Pi_ and k_-Pi_^BE^) or measured in this study (for V_ADP_ and V_ADP·Pi_), for reference. Each point is the result of the fit of the depolymerization rate curve extracted from at least 40 filaments. Black bars are means of replicates for a given condition. Indicated significances are from unpaired student t-tests. (ns : p>0.05 ; * : p≤0.05 ; ** : p≤0.01 ; *** : p≤10^-3^ ; **** : p≤10^-4^)

To test this hypothesis directly for *S. pombe*, we expressed *Sp*Act1 in a *Pichia pastoris* strain expressing SETD3, the enzyme responsible for the histidine 73 methylation in mammalian actins (Wilkinson et al., 2019). Using mass spectrometry, this was shown to produce actin that is 100% methylated on histidine 73 (Hatano et al., 2020). Similarly, using mass spectrometry, we validated that the vast majority of our SpAct1 recombinantly expressed in the presence of SETD3 was methylated on histidine 73 (Figure S8). We refer to this actin as *Sp*Act1^H73me^.

The resulting actin assembly dynamics for *Sp*Act1^H73me^ were then assessed in the same way as for *Sp*Act1 and *Sc*Act1. The kinetic rate constants we measured for ATP-actin at the barbed end, k_on_^ATP^ = 11.7±0.7 s^-1^.µM^-1^ and k_off_^ATP^ = 1.1±0.6 s^-1^, were indistinguishable from those of both *Sp*Act1 and *Oc*ACTA1 (Figure 4B, Table 3). The barbed end off-rate constant of ADP-actin, k_off_^ADP^ = 29.1±0.8 s^-1^, was also not significantly different from the one of *Sp*Act1 and was similarly accelerated compared to *Oc*ACTA1 (Figure 4B, Table 3). However, the *Sp*Act1^H73me^ barbed end had a roughly 2-fold higher affinity for Pi than its non-methylated counterpart, due to a faster binding and an unaffected unbinding of Pi at the barbed end (Figure 4C,D, Table 3). *Sp*Act1^H73me^ barbed ends also displayed a lower ADP·Pi-actin off-rate, k_off_^ADP·Pi^ = 2.8±0.1 s^-1^ (compared to 6.4±0.9 s^-1^ for non-methylated *Sp*Act1), which is still much faster than for *Oc*ACTA1 (Table3). The ionic and Hofmeister effects were as moderate as for its non-methylated counterpart (Figure S4C).

Interestingly, histidine 73 methylation reduces the Pi-release rate of *S. pombe* actin filaments approximately 6-fold: k_-Pi_ = 0.028±0.002 s^-1^ for *Sp*Act1^H73me^, compared to 0.16±0.06 s^-1^ for *Sp*Act1 (Figure 4D). Pi release for *Sp*Act1^H73me^ is still about 4-fold faster than for *Oc*ACTA1 (Jégou et al., 2011). Mammalian beta actin, which is closer to *Sp*Act1 in amino acid sequence identity than *Oc*ACTA1 (Figure S1), was reported to release Pi from filaments either at the same rate as *Oc*ACTA1 (Oosterheert et al., 2023) or 2.5-fold faster (Schahl et al., 2025) which would make it similar to *Sp*Act1^H73me^ in that respect. We found that Pi release from the barbed end was not significantly affected by histidine 73 methylation (Figure 4D).

Together, these results show that the absence of histidine 73 methylation accounts for most of the differences between *S. pombe* and mammalian actins in terms of Pi release from the filament, while it has little or no effect on the other rates we measured.

### Yeast actin filaments are slightly more flexible than mammalian actin filaments

The barbed end dissociation rate partly reflects the strength of the bonds between actin subunits within the filament, which has bearings on the mechanical stiffness of the filament. This made us wonder if *Sp*Act1 and *Sc*Act1 filaments were more flexible than *Oc*ACTA1 filaments. Previous literature indicates that *S. cerevisiae* actin filaments might be more flexible than their mammalian counterparts (Kang et al., 2012; X.-P. Xu et al., 2024; Stevenson et al., 2025; McCullough et al., 2011). We sought to confirm this and ask whether this property is conserved in yeast beyond *S. cerevisiae*. For that, we used a simple assay where we stick actin filaments to a coverslip coated with poly-L-lysine (Figure 5A) and determined their persistence Length Lp by fitting the average angular cosine correlation along filaments (Figure 5B-D, Methods). Interestingly, in our conditions, we found that *Sc*Act1 filaments, with Lp = 7.46±0.76 µm, were slightly but significantly more flexible than *Sp*Act1 filaments, for which Lp = 8.48±0.41 µm (Figure 5D). Both yeast actin filaments were more flexible than *Oc*ACTA1 filaments, for which we measured Lp = 9.59±0.59 µm. This decrease in persistence length was not rescued by the methylation of histidine 73, which did not seem to change the persistence length of *Sp*Act1 filaments (Figure 5D).

**Figure 5.**
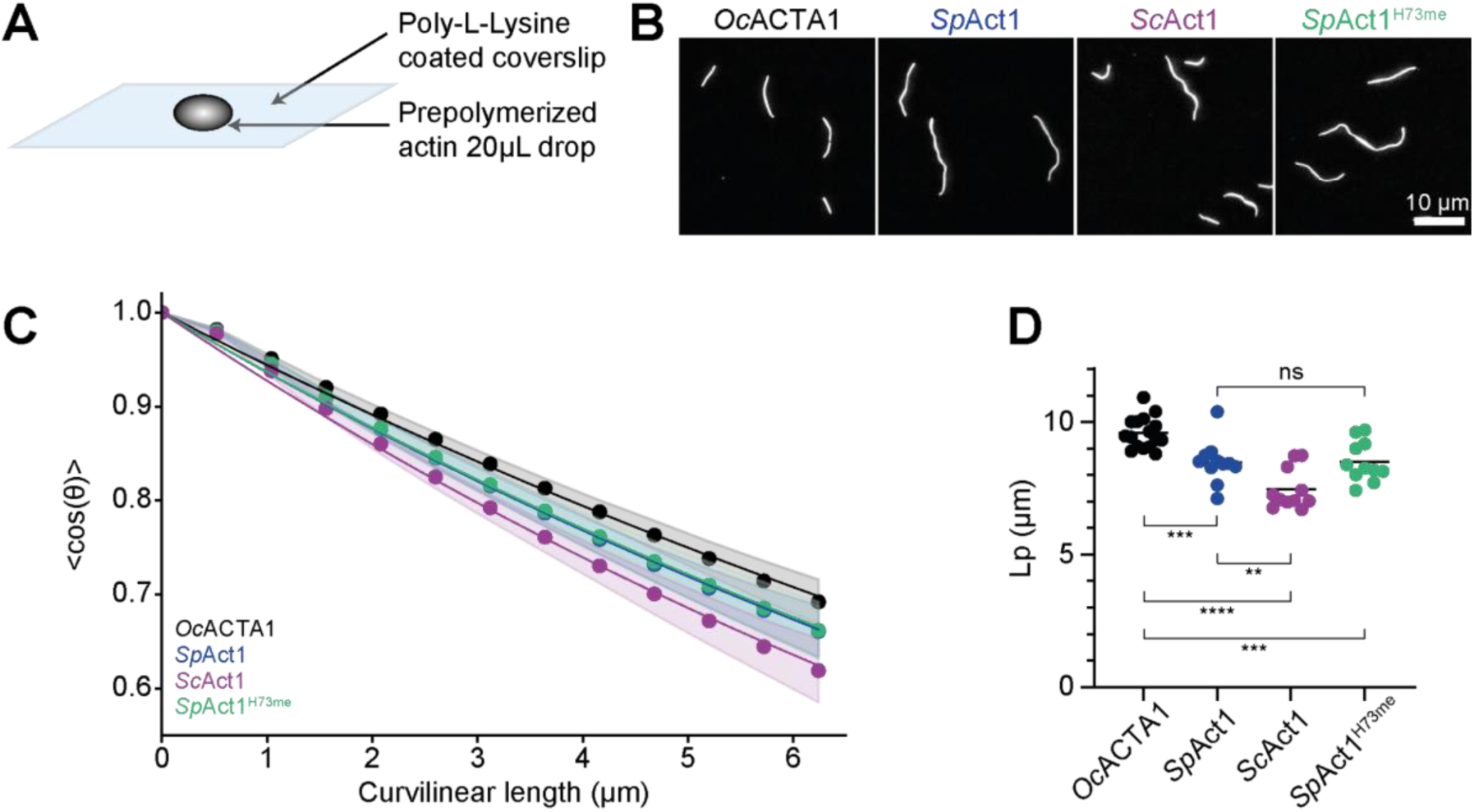
The persistence length of yeast actin filaments is reduced compared to rabbit actin independently of histidine 73 methylation. **A.** Method used to study the persistence length of actin filaments. Pre-polymerized ATP-ATTO488 F-actin is deposited onto a coverslip functionalized with poly-L-lysine. Snapshots of surface-attached filaments are quickly recorded by fluorescence microscopy before the solution dries. **B.** Example of a field of view using the method in A for *Oc*ACTA1 (black), *Sp*Act1 (blue), *Sc*Act1 (magenta) and *Sp*Act1^H73me^ (Green). **C.** Average cosine angle correlation as a function of the curvilinear distance between two points along actin filaments for the different actins. Data points for each actin are the mean values from at least 11 independent experiments, each containing at least 323 filaments. Shaded areas are standard deviations. Lines are exponential fits (see Methods). **D.** Plots of the persistence length for *Oc*ACTA1 (black), *Sp*Act1 (blue), *Sc*Act1 (magenta) and *Sp*Act1^H73me^ (Green). Each point is the result of the cosine correlation fit extracted from at least 323 filaments. Black bars are means of replicates for a given condition. Indicated significances are from unpaired student t-tests. (ns : p>0.05 ; ** : p≤0.01 ; *** : p≤10^-3^ ; **** : p≤10^-4^)

## Discussion

By directly comparing *Schizosaccharomyces pombe* (*S. pombe*) and *Saccharomyces cerevisiae* (*S. cerevisiae*) actin filaments, we showed that they have strikingly similar assembly dynamics, in spite of their evolutionary distance. The two actins differ mildly in their kinetics of disassembly, and not in a systematic fashion: for example, the off-rate is higher for *S. pombe* in ADP·Pi-actin, and higher for *S. cerevisiae* in ADP-actin (Table 3). These reaction rate constants should prove valuable for studies that aim to understand the differences between actin-driven mechanisms in the two yeasts, such as endocytosis (Nickaeen et al., 2022; Picco et al., 2025). The two actins differ also in filament stiffness, *S. cerevisiae* actin filaments having a lower persistence length. Differences in amino acid composition likely account for the kinetic and mechanical differences that we measured. For example, the W loop, that mediates longitudinal contacts between subunits in the filament, contains residues which differ in the two yeasts (shown in magenta in Figure 4A) and could explain in part why *S. cerevisiae* actin filaments are more flexible (Stevenson et al., 2025; Figure 5). In fact, this residue difference seems specific to *S. cerevisiae* among known eukaryotic actins (Figure S1) suggesting that increased flexibility may be a feature of *S. cerevisiae*.

Compared to metazoan actins, and in particular the widely studied alpha-skeletal actin from rabbit, we found that actin from the two yeasts assembled equally fast: the reaction rate constants for ATP-actin at the barbed end were indistinguishable at 25°C. The identical on-rate constant k_on_^ATP^ was not expected since it is not a necessary consequence of muscle ATP-actin barbed end elongation being diffusion-limited (Drenckhahn & Pollard, 1986) and the elongation rate of actin can be affected by a single point mutation (Greve et al., 2024). Furthermore, the identical off-rate constant k_off_^ATP^ indicates that the binding energy of the terminal subunit is very close to that of a muscle actin filament barbed end, in spite of the differences in the residues that mediate this binding (Figure 4A). *In vivo*, however, the actin filaments would not elongate at the same rate, in particular because of differences in temperature between these organisms. This suggests that the tremendous constraints that have kept these rates nearly identical over a billion years of evolution do not simply reflect the biological role of actin filaments in cells. Future studies should help determine the contributions of physiological and chemical factors in constraining these rates. For example, it would be interesting to know if purified actin filaments from other species also elongate at the same rate at 25°C. Experiments in that direction should benefit from the recent tools we have used, in particular the use of fluorescently labeled ATP to monitor the filaments. As already reported (McKane et al., 2006; Gressin et al., 2015), actin from yeast and actin from rabbit muscle can copolymerize to form hybrid filaments. Here, we find that these hybrid filaments elongate slower (Figure 1G-I, Figure S2), suggesting that the bonds between subunits from different species are weaker. However, in our experimental conditions, these mixed-species interfaces did not appear particularly fragile (Figure S10).

In contrast to elongation, we found that yeast actin filaments depolymerize markedly faster, both in ADP·Pi-and ADP-actin, compared to mammalian filaments. While depolymerization in the absence of actin monomers, as done in this study, is unlikely to occur in cells, these rate constants reveal fundamental differences between actins from the three species. Amino acid differences found in both yeast actins compared to mammals (residues marked in yellow in Figure 4A), could contribute to a weaker binding of the subunits, in particular isoleucine 43 (valine in mammals) which is located in the D-loop.

Yeast actin filaments also release inorganic phosphate (Pi) much faster than mammalian actin filaments, and this is largely due to the absence of methylation of histidine 73 in yeast (shown in green in Figure 4A). While mammalian alpha-skeletal and beta actins are both methylated on histidine 73, the observation that the latter releases Pi 2.5-fold faster (Schahl et al., 2025) suggests that other residues are involved in this process. A good candidate is methionine 16 (shown in pink in Figure 4A), found in the nucleotide-binding pocket of yeast actins and mammalian beta but not alpha-skeletal actin (leucine16, instead), and already suggested to favor Pi release based on structural data (Stevenson et al., 2025). The importance of histidine 73 methylation highlights the need to control key post-translational modifications *in vitro*, and one should note that recombinant actins expressed in insect cells are likely to be artificially histidine 73-methylated. In *S. pombe* actin, the methylation of histidine 73 increases the affinity of Pi for the actin subunit at the barbed end (Figure 4C) by increasing the on-rate constant of Pi into the barbed end subunit (k_+Pi_^BE^, Table 3) without affecting its off-rate constant (k_-Pi_^BE^, Figure 4D, Table 3). Histidine 73 methylation also slows down the depolymerization of ADP·Pi-actin (Figure 4C, Table 3).

The release of Pi from the barbed end is also faster in yeast than in mammals, but to a lesser extent than the release of Pi from the filament core, and this effect does not seem related to the absence of histidine 73 methylation in yeast (Figure 4D, Table 3). Instead, it is likely due to larger fluctuations of the terminal subunit in yeast, which may ease the opening of the Pi escape route (the so-called “back door”; Oosterheert et al., 2023; Wriggers & Schulten, 1999) or allow another route through which the Pi can escape. In the monomeric form, both yeast actins have been reported to exchange their nucleotide faster than mammalian actins (Chen et al., 1993; Ti & Pollard, 2011), and our observations using ATP-ATTO488 are consistent with this (Fig S7).

The cytoskeleton is particularly interesting to study in the light of evolution because it serves essential functions in diverse organisms across the tree of life. For example, actin filaments purified from apicomplexans and from *Leishmania*, which are evolutionary very far from metazoans, appear to depolymerize rapidly *in vitro* (Lu et al., 2019; Kotila et al., 2022; Hvorecny et al., 2024). This is commonly interpreted as a means for these organisms to have actin cytoskeletons that turn over very fast, in spite of having a limited set of regulatory proteins to accelerate their disassembly. *S. pombe* and *S. cerevisiae*, which are closer to metazoans, provide a somewhat intermediate situation allowing us to test and expand this interpretation. They also have very dynamic actin cytoskeletons but they have orthologs of the key ABPs found in metazoans (Moseley & Goode, 2006). The rapid depolymerization rates we report here, compared to mammalian actin filaments, could be a means for yeasts to operate efficiently at lower temperatures. The rapid Pi-release rates from the filament core and barbed ends that we report, as well as the lower affinity of Pi for the barbed ends, could be important for yeasts to operate with high cytosolic concentrations of Pi (van Eunen et al., 2010). Also, since yeasts have fewer different actin networks than animal cells, they may not need such a wide range of turnover rates and may thus benefit from having less stable actin filaments. The elongation of actin filaments can be regulated by several factors in yeast cells (Billault-Chaumartin et al., 2022; Gonzalez Rodriguez et al., 2023; Homa et al., 2021; Scott et al., 2011). Nonetheless, the rates we report here indicate that, with an estimated 4 µM of actin monomers in the cytosol of *S. cerevisiae* (Gonzalez Rodriguez et al., 2023), actin filaments on their own would elongate at roughly 100 nm/s at 25°C, which appears compatible with the fast internalization observed at endocytic sites, for example (Kaksonen et al., 2005).

Yeast, and in particular *S. pombe* and *S. cerevisiae*, are powerful model organisms to study the actin cytoskeleton. Conveniently, they possess a few, well-defined and well-characterized actin networks (Kovar et al., 2011; Mishra et al., 2014), which can be studied to derive universal properties of the actin cytoskeleton. As already highlighted years ago (Berro et al., 2010; Pollard & Berro, 2009; Pollard & De La Cruz, 2013), taking full advantage of these systems requires knowing the underlying reaction rates. Here, we provide a number of essential rates that were missing (Figure S3, Table 3) and should contribute to improving our understanding of actin network dynamics in yeasts. Nonetheless, several important reaction rates remain unknown, regarding the interactions of actin filaments and monomers with ABPs in yeast. Since actin-ABP interactions can be very species-specific (Ghasemi et al., 2024; Kotila et al., 2022), future *in vitro* studies will have to carefully match the origins of the proteins in order to reliably decipher the regulation of actin dynamics.

## Declaration of interests

The authors declare no competing interests.

## Author contributions

IBC, GRL and AJ conceived the project. HW performed and quantified part of the experiments in Figure S5 and developed the code used to quantify experiments relative to Pi-release rate. IBC performed all other experiments and quantifications. AG purified the native yeast actin used in Figure S5. HW purified the mammalian actins used in Figure S5. IBC purified all other proteins. AM, AJ and GRL acquired funding. IBC and GRL wrote the first draft of the manuscript, which was revised by all authors.

## Acknowledgments

We thank Mohan Balasubramanian for *P. pastoris* strains and plasmids, the Protéoseine facility for mass spectrometry analysis and Sophie Martin for comments on the manuscript. We acknowledge funding from Fondation pour la Recherche Médicale (grant EQU202203014630 to G.R.L.), from the Fondation Bettencourt-Schueller (grant Impulscience to A.J.), from the National Research Foundation (NRF) Singapore under its Mid-Sized Grant (MSG) (NRF-MSG-2023-0001 to A.M.) and from the National University of Singapore through the Mechanobiology Institute (A-0003467-00-00 to A.M.).

## Supplementary figures

**Figure S1 – relative to Figures 1 and 4.**
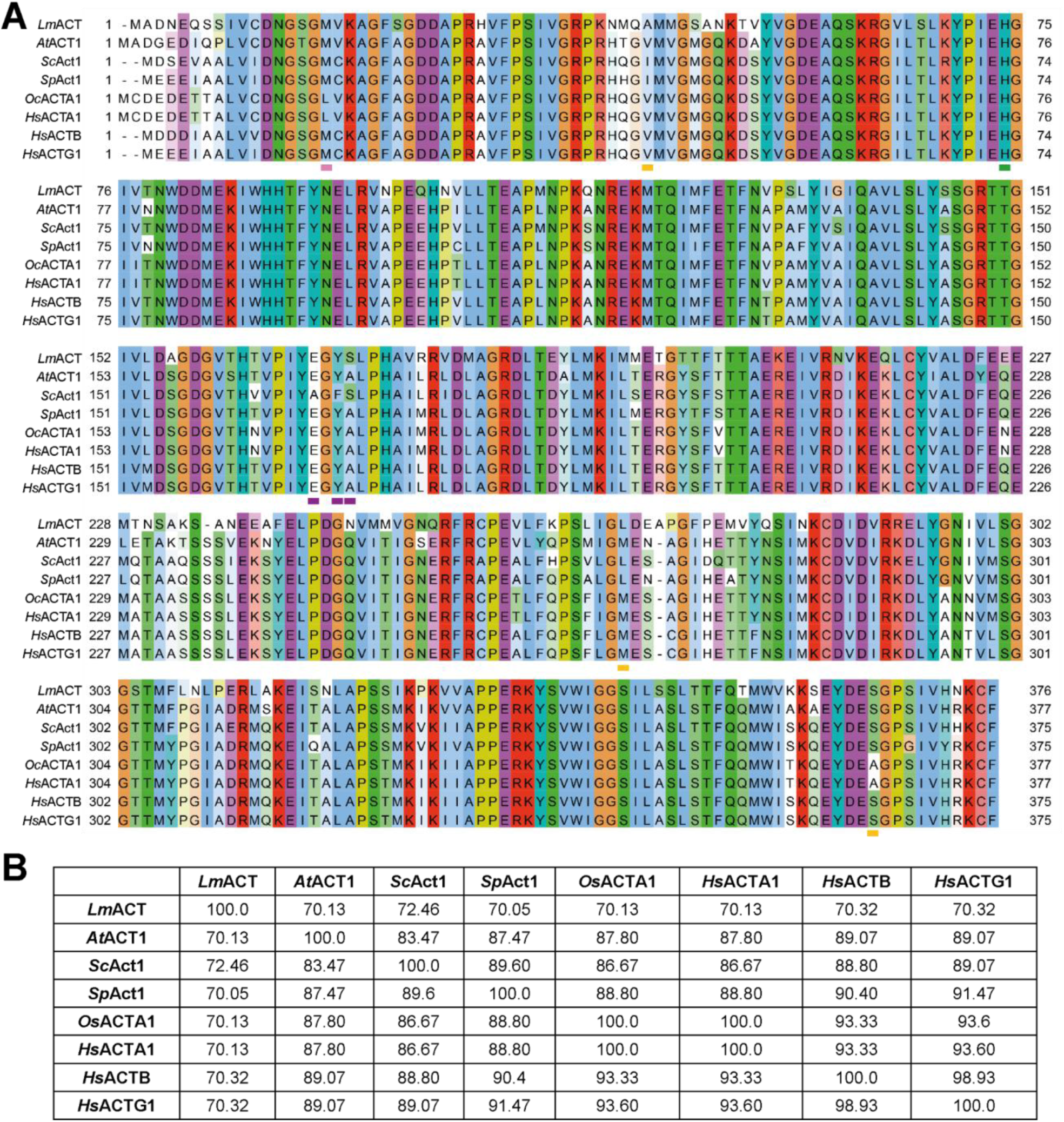
Selected actins proteic sequence alignment. **A.** ClustalO alignment of actin sequences from *Lm* = *Leishmania major* (Q9U1E8), *At* = *Arabidopsis thaliana* (P0CJ46), *Sc* = *Saccharomyces cerevisiae* (P60010), *Sp* = *Schizosaccharomyces pombe* (P10989), *Oc* = *Oryctolagus cuniculus* (alpha skeletal muscle, P68135), *Hs* = *Homo sapiens* (ACTA1 = alpha skeletal muscle, P68133 ; ACTB = beta, P60709 ; ACTG2 = gamma cytoplasmic, P63261). The pink box indicates *Sc* M16 residue highlighted in Figure 4A which is conserved in all depicted species but alpha skeletal muscle actins. The yellow boxes indicate *Sc* I43, L269 and S324 residues highlighted in Figure 4A which are divergent in both *Sc* and *Sp* relative to almost all other species. The green box indicates residue H73 highlighted in Figure 4A, methylated in mammalian actins but unmethylated in *Sc* and *Sp* (Table 2). The magenta boxes indicate *Sc* A167, F169 and S170 residues, highlighted in Figure 4A, which are highly specific to *Sc*. A phylogenetic tree from a subset of the actins from this alignment is displayed in Figure 1A. **B.** Percent identity matrix of the aligned actin sequences shown in A.

**Figure S2 – relative to Figure 1.**
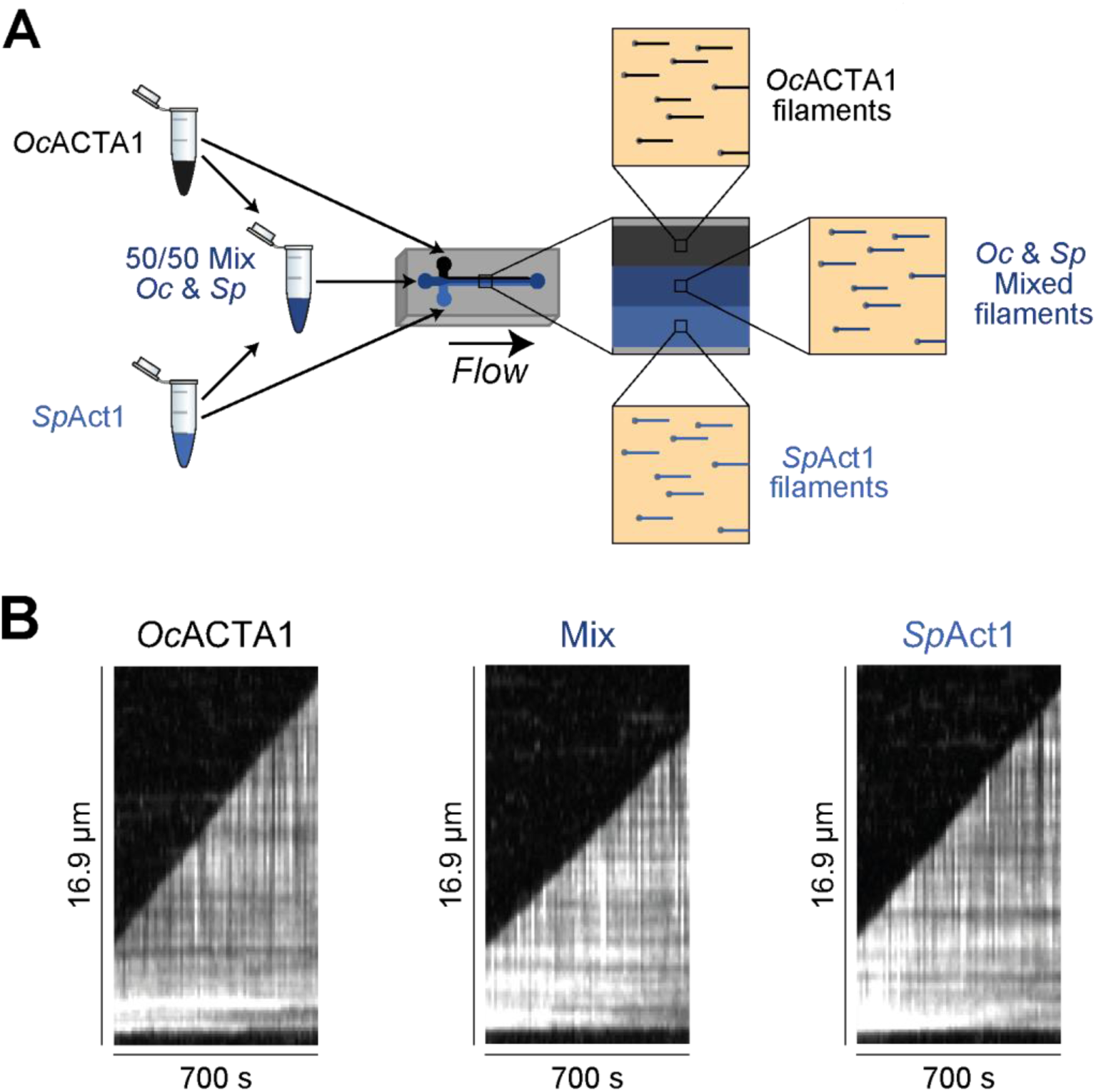
Actins from different species are relatively incompatible. **A.** Method used to study the compatibility of actins from different species. Pure actin solutions of *Oc*ACTA1, *Sp*Act1 and *Sc*Act1, labelled with a half molar ratio of ATP-ATTO488, are prepared and used to obtain 50/50 mixes containing two actin types. For one given experiment, two pure solutions in F-buffer (yellow) and the resulting mix are simultaneously injected in three different entry channels of a microfluidics chamber and their elongation rate is recorded and averaged over 30 filaments. **B.** Example of Kymographs of an actin filament over the course of an experiment as in A, using 0.5 µM *Oc*ACTA1, 0.5 µM *Sp*Act1, or a 50/50 mix of both actins, in all cases with 0.3 µM of ATP-ATTO488 in F-buffer.

**Figure S3 – relative to Table 3.**
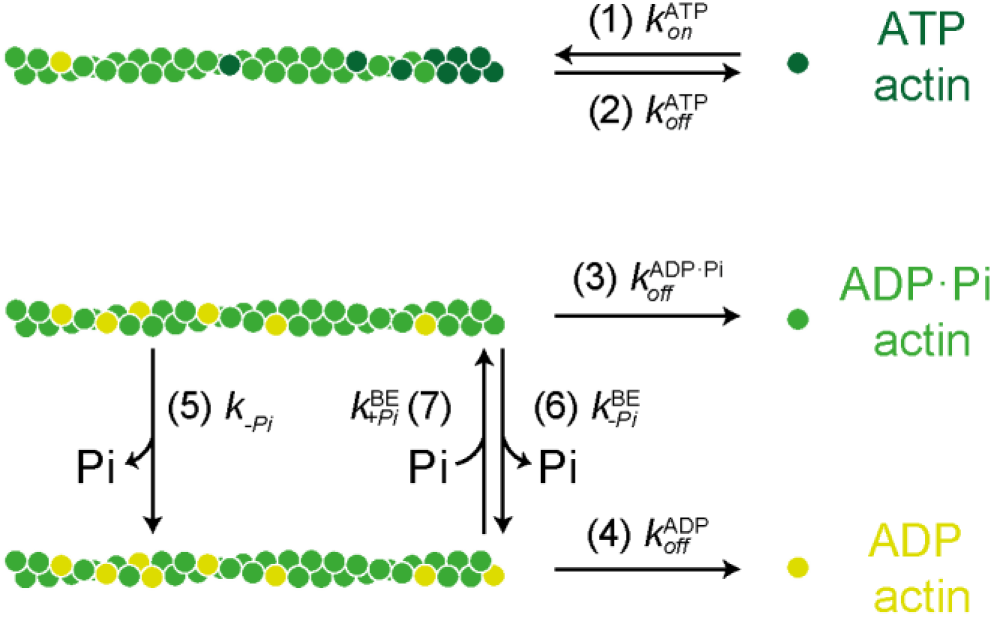
Schematic representation of the biochemical pathways characterized in this study. **(1)** Association rate constant of an ATP-actin monomer onto an ATP-actin barbed end. **(2)** Dissociation rate constant of an ATP-actin monomer from an ATP-barbed end. **(3).** Dissociation rate constant of an ATP·Pi-actin monomer from an ATP·Pi-barbed end. **(4)** Dissociation rate constant of an ADP-actin monomer from an ADP-barbed end. **(5)** Dissociation rate constant of an inorganic phosphate molecule from an actin subunit located inside of a filament. **(6)** Dissociation rate constant of an inorganic phosphate molecule from an actin subunit located at the barbed end. **(7)** Association rate constant of an inorganic phosphate molecule binding to an actin subunit located at the barbed end. Note that reactions (3), (4) and (5) are reversible, but the rate constants of their reverse reactions have not been measured in this study.

**Figure S4 – relative to Figures 2 and 4.**
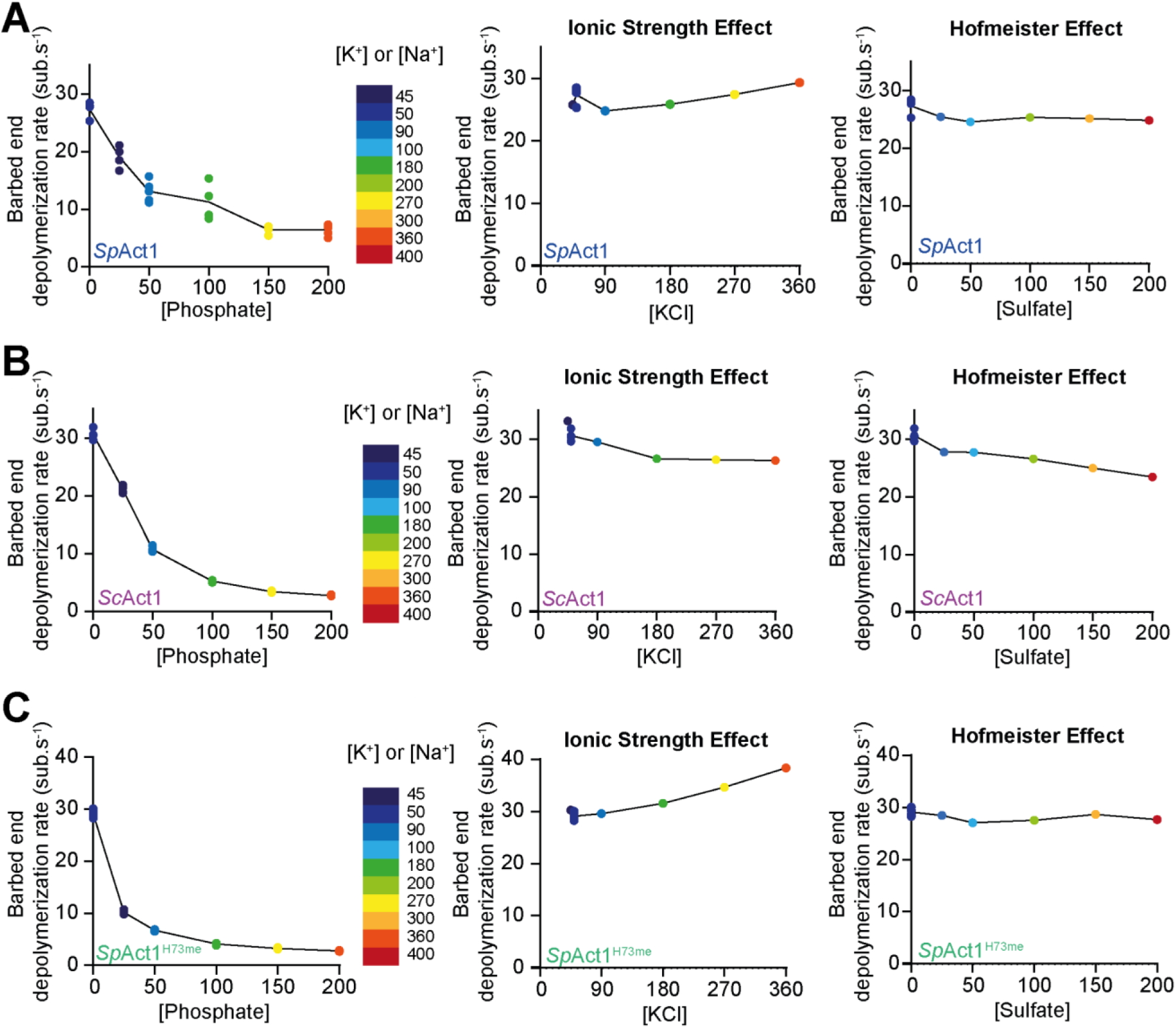
Estimation of the impact of the ionic strength and Hofmeister effects of inorganic phosphate on the depolymerization of *Saccharomyces cerevisiae* and *Schizosaccharomyces pombe* actin filaments. **A-C.** Plots of the barbed end depolymerization rate, color coded as a function of the positive ion concentration, as a function of (Left) the phosphate concentration as in Figures 2C and 4C, (Middle) the KCl concentration and (Right) the sulfate concentration for *Sp*Act1 (A, blue), *Sc*Act1 (B, Magenta) and *Sp*Act1^H73me^ (C, Green). Each point is the mean of a replicate where the elongation rate of at least 15 filaments was quantified.

**Figure S5 - relative to Figure 1.**
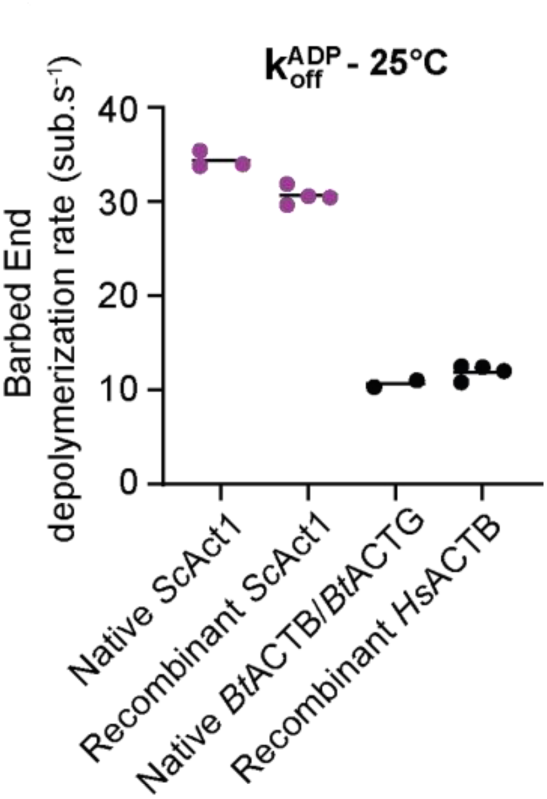
Using recombinant rather than native actin does not significantly alter ADP-actin depolymerization rate. Plot of the ATP-actin depolymerization rate for native and recombinant *S. cerevisiae* actin (magenta) and mammalian actin (black). Data for recombinant *Sc*Act1 is a duplicate from figure 1F included for comparison. Note that due to availability, for recombinant mammalian actin, *Homo sapiens* beta cytoplasmic actin was used, both N-terminally acetylated and histidine 73 methylated as it is *in vivo*, and for native mammalian actin, *Bos taurus* thymus actin was used, which has been shown to be a mix of both beta and gamma cytoskeletal actin isoforms (Vandekerckhove & Weber, 1978) as it is a better reference for the aforementioned native actin than alpha skeletal muscle rabbit actin. Each point is the mean of an independent replicate where the depolymerization rate of 30 filaments was quantified, and black bars are means of the replicates for a given condition.

**Figure S6 - relative to Figure 1 and 4.**
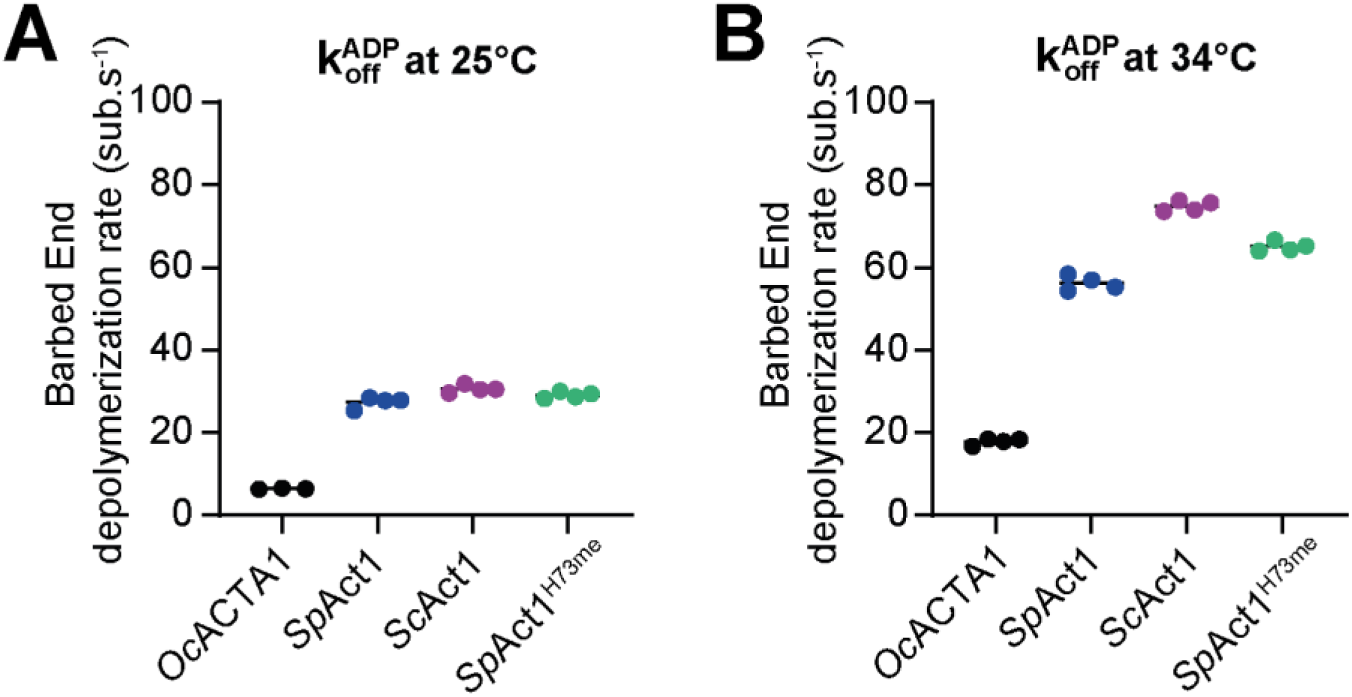
Temperature dramatically accelerates ADP-actin depolymerization rate in all actins considered. **A.** Plot of the barbed end depolymerization rate k_off_^ADP^ for *Sp*Act1 (blue), *Sc*Act1 (magenta), *Sp*Act1^H73me^ (Green) and *Oc*ACTA1 (black) at 25°C. These values are the ones shown in Figures 2D and 4B, at 0 µM actin, obtained after polymerizing the filaments, and are shown here to ease comparison with panel B. Each point is the mean of an independent replicate where the depolymerization rate of 30 filaments was quantified, and black bars are means of the replicates for a given condition. Dotted line is the value obtained for *Oc*ACTA1 in Wioland et al., 2019, for reference. **B.** Plot of the barbed end depolymerization rate k_off_^ADP^ for *Oc*ACTA1 (black), *Sp*Act1 (blue), *Sc*Act1 (magenta) and *Sp*Act1^H73me^ (Green) at 34°C. Each point is the mean of an independent replicate where the depolymerization rate of 30 filaments was quantified, and black bars are means of the replicates for a given condition.

**Figure S7.**
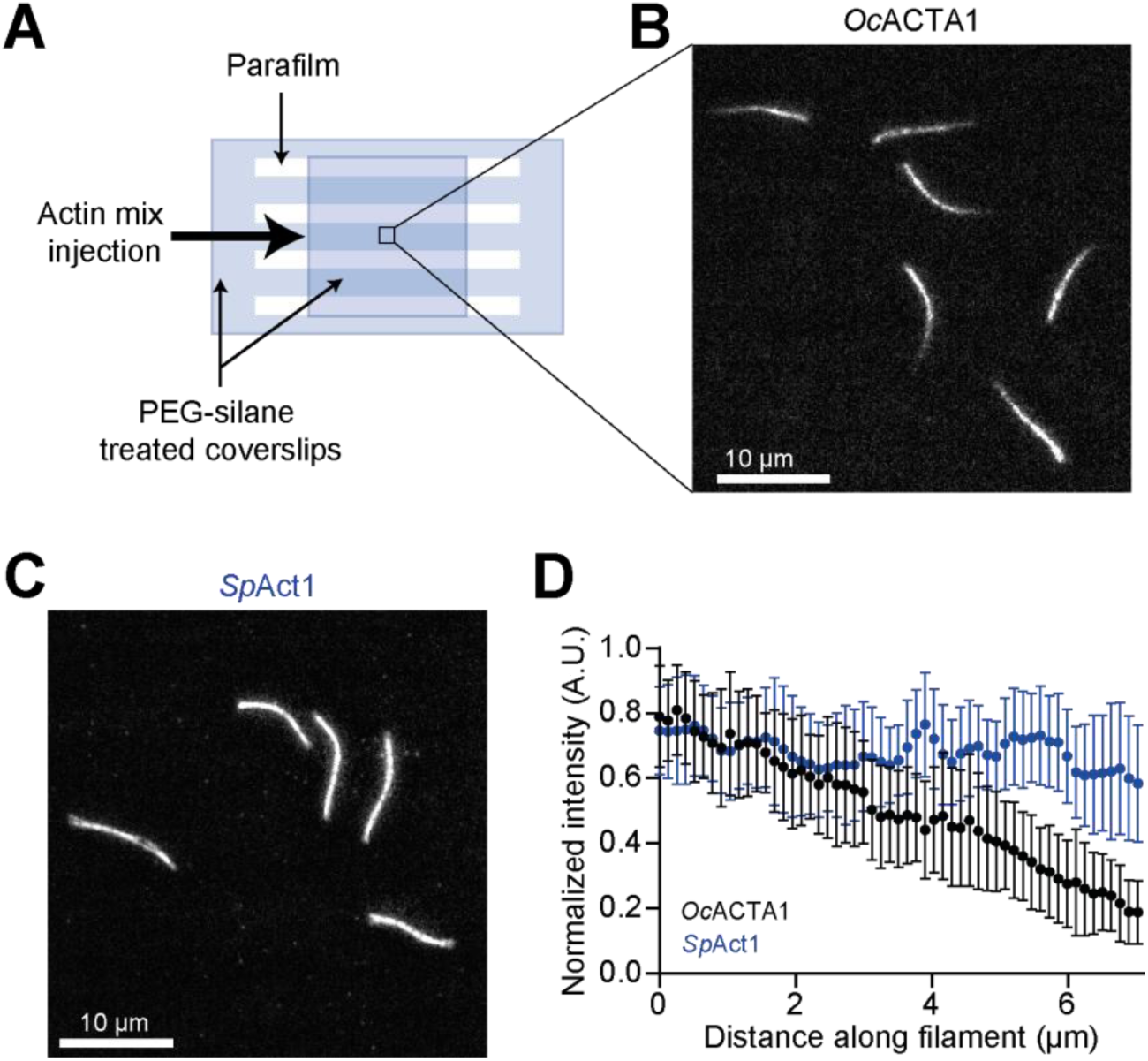
The nucleotide exchange is faster in *Schizosaccharomyces pombe* than in rabbit monomeric actin. **A.** Method used to study the exchange rate of the nucleotide inside of monomeric actin. G-Actin was mixed with ATP-ATTO488 and immediately injected in polymerizing conditions into an open chamber canal made with coverslips and parafilm strips. Filaments were maintained close to the surface by using methylcellulose. Movies of the polymerizing filaments were acquired by TIRF microscopy. If the exchange inside of the monomer is slow and polymerization fast, the labelling fraction will increase over time and hence the intensity along the filament will increase from pointed end to barbed end. Instead, if the exchange is fast, the intensity will be homogenous. **B.** Example of a field of view using the method as in A with 0.35 µM *Oc*ACTA1 and 0.175 µM ATP-ATTO488 in F-buffer without ATP supplemented with 0.18% methylcellulose. **C.** Example of a field of view using the method as in A with 0.35 µM *Sp*Act1 and, 0.175 µM ATP-ATTO488 in F-buffer without ATP supplemented with 0.18% methylcellulose. **D.** Plot of the normalized mean intensity profile (see Methods) of 20 filaments from an experiment as in A for *Oc*ACTA1 (black) and as in B for *Sp*Act1 (blue). Filaments were registered by their brighter end, defined as the origin on the graph. Error bars are standard deviations.

**Figure S8.**
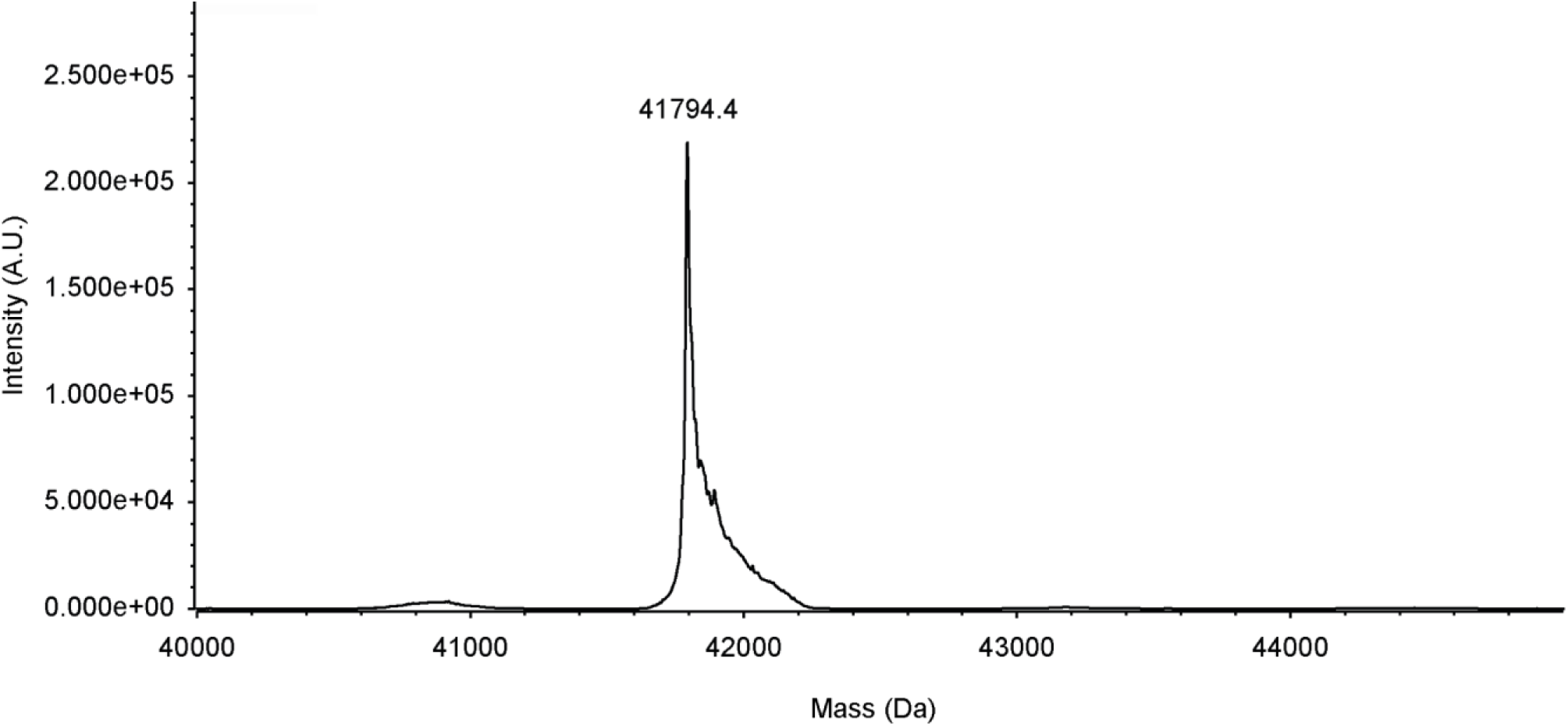
The vast majority of *Schizosaccharomyces pombe* actin recombinantly expressed in *Pichia pastoris* in the presence of SETD3 is methylated. Deconvoluted LC-MS spectra of intact *Sp*Act1^H73me^. Expected sizes are 41738.66 Da for unmodified SpAct1(Y371H) , 42.01 Da for acetylation, 14.02 Da for methylation, i.e. 41794.69 Da for an actin both acetylated and methylated which is consistent with the weight of the main peak. Note that there is no peak at 41780.67 Da, the expected mass of unmethylated actin.

**Figure S9 - relative to Figure 3.**
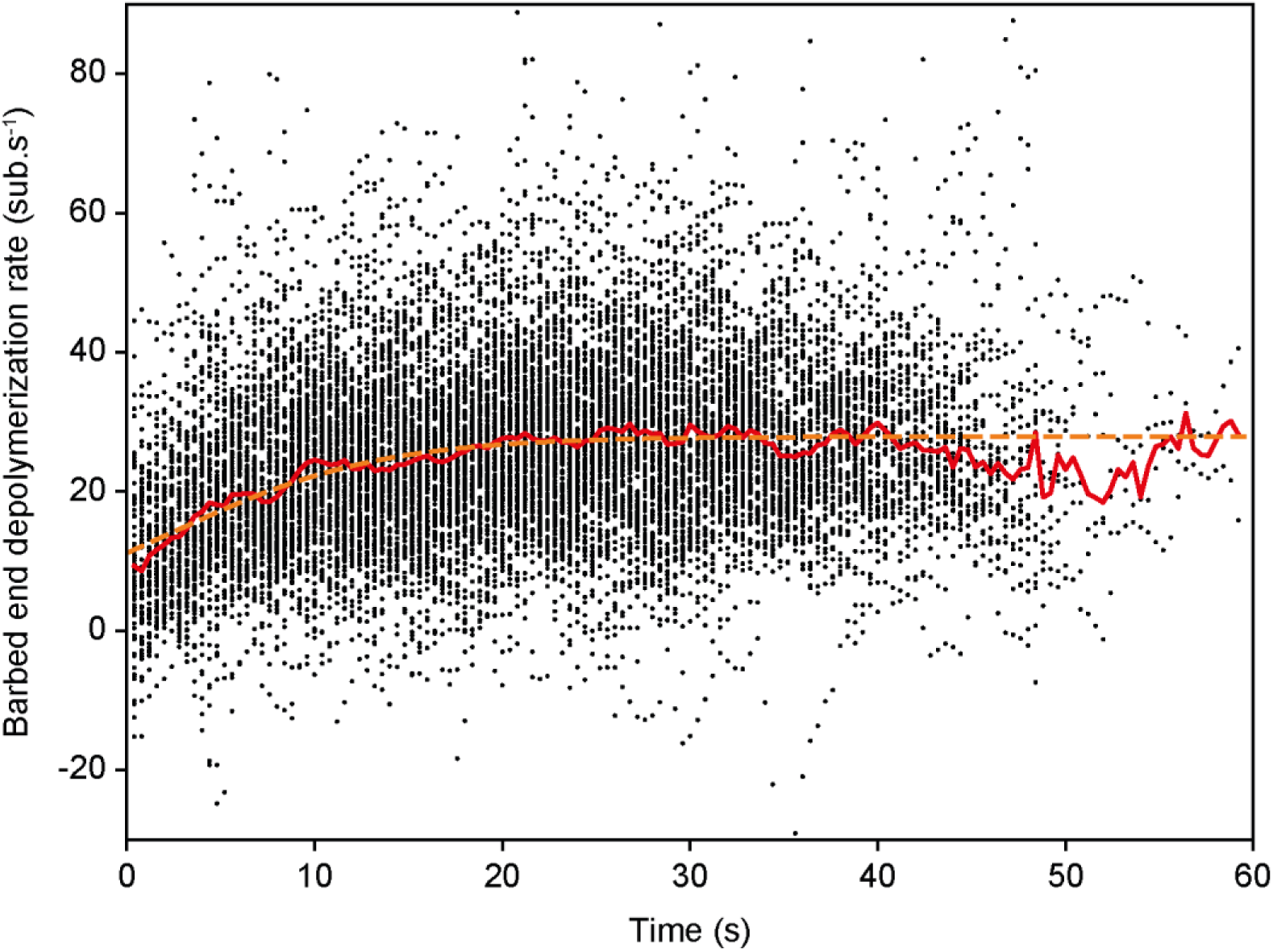
Fitting depolymerization data in Pi-release experiment. Example of a fit of the depolymerization (see Methods) for experiments such as the one shown in Figure 3A-B, where filaments were elongated using 0.8 µM *Sp*Act1, 0.4 µM ATP-atto488 and 50 mM phosphate prior to their disassembly in F-buffer. The depolymerizing barbed ends of 183 filaments were tracked over time using a custom Python script (Kotila et al., 2022; Schahl et al., 2025). Each black dot is the instantaneous depolymerization rate extracted by a linear fit over 5 frames, centered on time t, for a single filament. The red line is the average over the filament population. The orange dashed line is the fit of this average, using the following equation:

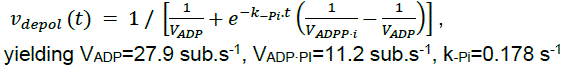

**Figure S10 – relative to Figure 1.**
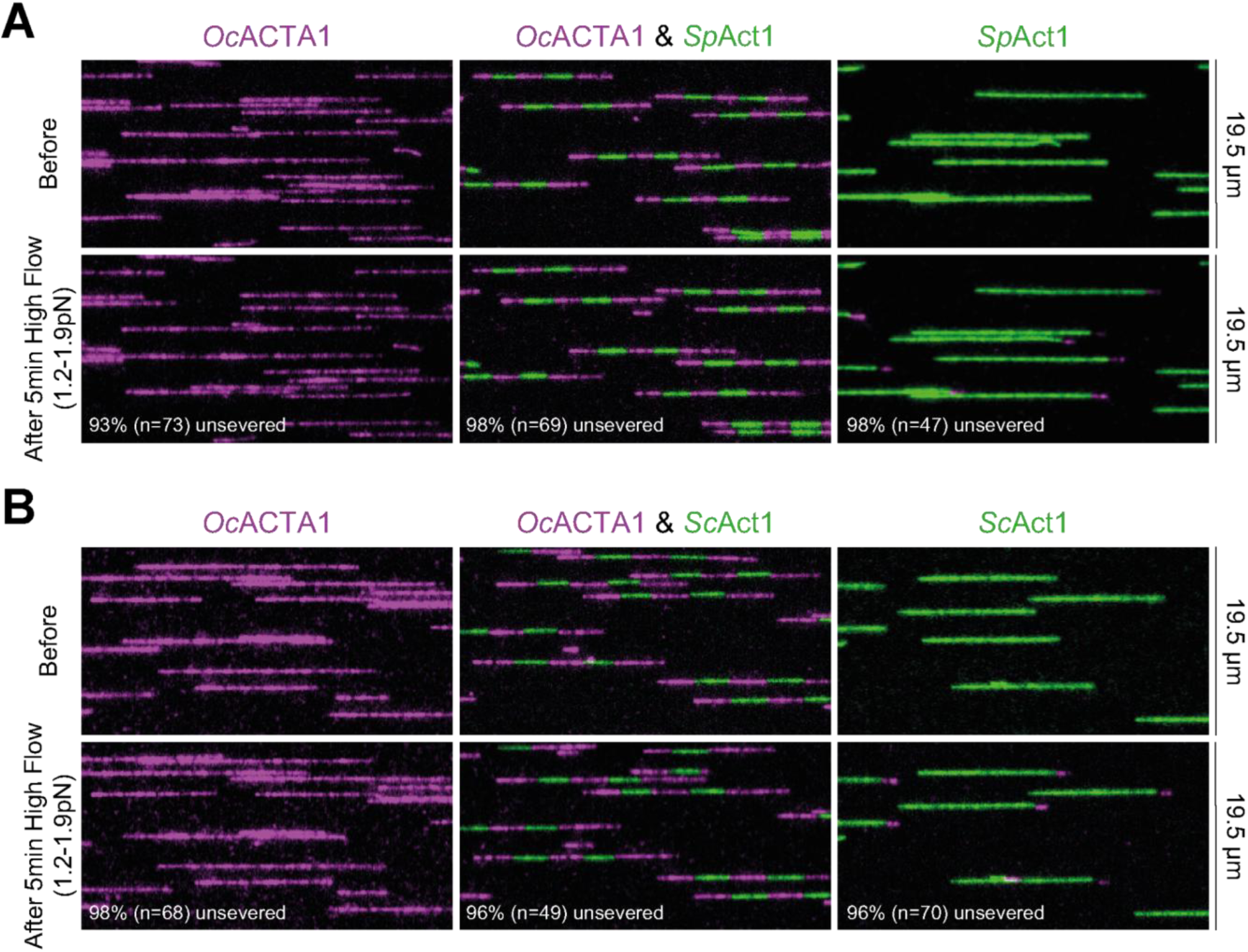
Boundaries between actins from different species are not particularly fragile **A-B.** Typical images from a microfluidics experiment where filaments were elongated using 0.6 µM *Oc*ACTA1 (10% labelled on lysines with Alexa568, in magenta, images on the left), or 0.6 µM yeast actin (*Sp*Act1 (A) or *Sc*Act1 (B), labelled with 0.3 µM ATP-atto488, in green, images on the right), or using actin from both channels in alternance (producing striped filaments, images in the middle). The three types of filaments were produced in different regions of the microfluidics chamber, and were simultaneously subjected to a fast-flowing solution of 0.2 µM *Oc*ACTA1 (10% labelled on lysines with Alexa568, in magenta) to compare their stability. After 5min of high flow (average tension within 1.2-1.9 pN), 93% (n = 73) of fully *Oc*ACTA1 filaments, 98% (n = 69) of striped filaments and 98% (n = 47) of fully *Sp*Act1 filaments remain unsevered in the Oc/Sp experiment and 98% (n = 68) of fully OcACTA1 filaments, 96% (n = 49) of striped filament and 96% (n = 70) of fully *Sc*Act1 remain unsevered in the Oc/Sc experiment.

## Supplementary legends

**Movie S1 – relative to Figure 1**. Polymerization of *Sp*Act1 filaments.

TIRF imaging of an experiment as in Figure 1B using 0.5 µM *Sp*Act1 and 0.25 µM ATP-ATTO488 (cropped field of view). A snapshot of this movie is displayed in Figure 1C, and the kymograph of a single filament is displayed in Figure 1D. The time stamp in min:s is displayed at the top left corner and the scale bar at the bottom right corner.

**Movie S2 – relative to Figure 2**. Depolymerization of *Sp*Act1 filaments in 25 mM phosphate buffer.

TIRF imaging of an experiment as in Figure 2A using 1 µM *Sp*Act1, 0.5 µM ATP-ATTO488, and 25 mM phosphate (cropped field of view). A kymograph of a single filament is displayed in Figure 2B. Please note that *Sp*Act1 and ATP-ATTO488 are only flowed in during the polymerization step, of which we only show the final moments before switching to depolymerization. The condition is displayed at the top right corner of the movie (“P25” refers to the buffer, which contains 25 mM phosphate). The time stamp in min:s is displayed at the top left corner and the scale bar at the bottom right corner.

## Methods

### Proteins

*Oc*ACTA1 (Figures 1, 2, 5, S2, S6, S7, S10)

Rabbit alpha skeletal muscle native actin was purified from rabbit muscle acetonic powder as previously published (Spudich & Watt, 1971).

Briefly, rabbit muscle acetonic powder was obtained from fresh rabbit muscles which were chopped then washed by 2 rounds of resuspension in Extraction Buffer (500 mM KCl, 50 mM KHCO3) and centrifugation (4000 g, 10 min., 4 °C), then two rounds of resuspension in water whose pH was adjusted with NaCO_3_ to 8.6 and centrifugation (4000 g, 10 min., 4 °C). The final pellets were resuspended by manual agitation, frozen and blended in −20°C acetone. The blended frozen muscles were filtered and the operation was repeated twice, and the resulting muscle paste was spread out and dried overnight. The powder was recovered and stored at −20°C until use.

Acetonic powder was resuspended in X Buffer (2 mM Tris pH 7.8, 0.5 mM ATP, 0.1 mM CaCl_2_, 1 mM DTT, 0.01 % NaN_3_) and centrifuged (40000 g, 45 min., 4 °C). The supernatant was collected and filtered through glasswool. KCl was added to a final concentration of 3.3 M to remove contaminants and the solution centrifuged and filtered as before. The supernatant was then dialyzed O/N against 32 supernatant volumes of Dialysis Buffer (2 mM Tris pH 7.8, 1 mM MgCl_2_, 1 mM DTT), which brought the KCl concentration to 0.1 M. KCl was then added to a final concentration of 0.8 M and the solution was incubated under agitation for 1h30 at 4°C, then ultracentrifuged (100000 g, 3h30, 4 °C). The resulting pellet was resuspended with a potter in X buffer supplemented with 40 mM KCl and 2 mM MgCl_2_ and incubated overnight at 4 °C. KCl was then added to increase its concentration to 0.8 M and the solution was further incubated under agitation 1h30 at 4 °C then ultracentrifuged (100000 g, 3h30, 4 °C). Repeating those steps as described here allows for a better detachment of actin binding proteins. The resulting pellet was resuspended with a potter in G-buffer (2 mM Tris pH 7.8, 0.1 mM CaCl_2_, 0.01 % NaN_3_, 0.2 mM ATP, 1 mM DTT) and dialysed against the same buffer to depolymerize filaments at 4 °C for 3 days. The solution was recovered, ultracentrifuged (400000 g, 45 min., 4°C) and injected into an equilibrated Superdex 200 Hiload column (GE Healthcare) and actin was eluted with G-buffer into 1 mL fractions. Actin-containing fractions were identified by absorbance at 290 nm, validated by electrophoresis and pooled. The left-most fractions were excluded as they might contain actin dimers. The concentration was determined by measuring the OD at 290 nm. Actin was stored on ice for up to 8 weeks or flash frozen into liquid nitrogen and placed at −80 °C for long term storage.

The actin used in Figure S10 was fluorescently labelled on F-actin surface accessible lysines with Alexa568 succinimidyl ester (Life Technologies) as per manufacturer recommendation. Briefly, the aforementioned native alpha skeletal muscle actin was dialysed overnight in modified F-buffer (20 mM PIPES pH 6.9, 100 mM KCl, 0.2 mM ATP, 0.2 mM CaCl_2_) for polymerization. The resulting filaments were incubated with a 5x excess of fluorophore for 2h at room temperature on a rotating wheel, then ultracentrifuged (350,000 g, 30 min., Room Temperature (RT)). The resulting pellet was resuspended in G-buffer with a potter and the actin was left to depolymerize on ice for 2 hours. A new round of polymerization was induced by adding 400 mM KCl and 2 mM MgCl2 for 1 hour at RT. The filaments were ultracentrifuged (350,000 g, 30 min., RT) and resuspended in G-buffer with a potter. The solution was dialysed overnight against G-buffer, then ultracentrifuged (350,000 g, 30 min., 4 °C). The concentration and labelling fraction were determined by measuring the OD at 280 and 578nm. The final labelled actin solution was stored on ice in G-buffer for up to 8 weeks. A 10% labelling fraction was used for experiments in Figures S10 and obtained by diluting with unlabelled actin.

*Bt*ACTB/*Bt*ACTG (Figure S5)

*Bos taurus* thymus native actin was purified from calf thymus using a gelsolin affinity method adapted from Funk et al., 2019.

The MBP-6xHis-G4-G6 mouse Gesolin fragment was produced recombinantly in Rosetta2 competent cells. The pRSF plasmid containing the fragment (addgene #188454) was a kind gift from Pr. Rodriguez (Ceron et al., 2022). Expression of a 1 L TB culture of an exponentially growing positive recombinant was induced with 1mM IPTG for 16 h O/N. The cells were pelleted at 9000g for 20min, washed with PBS, centrifuged again and flash frozen in liquid nitrogen and stored at −80°C until use, when the pellet was thawed in 50mL Lysis Buffer (10 mM Tris pH 8.0, Imidazole pH 8.0, 500 mM NaCl, 5 mM CaCl_2_, 5 mM β-mercaptoethanol, 1 for 150 mL Complete EDTA-free protease inhibitor tablet, 10 mg/mL lysozyme), sonicated on ice for 5 minutes (10 s on, 20 % off, 100 % amplitude) and centrifuged at 100,000 g for 30 min at 4 °C. The supernatant was recovered, filtered through a 0.22 µm membrane and injected onto a 2 x 5 mL HisTrap stacked column (Cytiva), pre-equilibrated with 50 mL Equilibration Buffer (10 mM Tris pH 8.0, 500 mM Imidazole) and 50 mL Wash Buffer (10 mM Tris pH 8.0, 50 mM KCl, 20 mM Imidazole, 5 mM CaCl_2_, 0.2 mM ATP pH 8.0, 5 mM β-mercaptoethanol). The column was washed with 50 mL Wash Buffer and kept at 4 °C until the thymus extract is ready.

Fresh calf thymus from the butcher was blended into HOLO Buffer (10 mM Tris pH 8.0, 20 mM Imidazole pH 8.0, 5 mM CaCl_2_, 1 mM ATP pH 8.0, 5 mM β-mercaptoethanol, 1 mM PMSF, 1 for 200 mL EDTA-free Protease cocktail inhibitor tablets (Sigma)) and supplemented with 2.5 mM β-mercaptoethanol after homogenizing, then the pH was adjusted to 8.0 with Tris. The resulting mixture was centrifuged at 9000 rpm (F14-6×250y rotor) for 45 min at 4 °C. The supernatant was filtered through glasswool and the eluate was centrifuged again at 35000 rpm (Ti45 rotor) for 1 h at 4 °C. The supernatant was recovered, supplemented with 50mM KCl and 20mM Imidazole pH8.0 and filtered through 0.45 µm membranes. The actin-containing eluate was then injected onto the Gelsolin decorated HisTrap column. The column was then washed with 10 CV of Wash buffer (10 mM Tris pH 8.0, 20 mM Imidazole pH 8.0, 50 mM KCl, 5 mM CaCl_2_, 0.2 mM ATP pH 8.0, 5 mM β-mercaptoethanol). The actin was eluted through a gradient from Wash Buffer to Actin Elution Buffer (10 mM Tris pH 8.0, 20 mM Imidazole pH 8.0, 50 mM KCl, 5mM EGTA pH 8.0, 0.2 mM ATP pH 8.0, 5 mM β-mercaptoethanol) into 1 mL fractions into a 96 well plate where each well was prefilled with 20 µL 1M MgCl_2_. Pertinent fractions were identified as peak of fluorescence at 290 nm and further validated by electrophoresis, left to polymerize at RT for 3-4h, then pooled. The resulting solution was supplemented with 100 mM KCl and ATP concentration was adjusted to 0.5 mM, and polymerization was allowed to continue for 2 h at RT. The solution was then ultracentrifuged at 80000 rpm (TLA 100.3 rotor) at RT for 30 min. The resulting pellet was resuspended with a potter into G-buffer (10 mM Tris pH 8.0, 0.1 mM CaCl_2_, 0.01 % NaN_3_, 0.1 mM ATP, 1 mM DTT) and dialysed against the same buffer to depolymerize actin filaments. Depolymerization was aided by sonication. The solution was then ultracentrifuged at 90,000 rpm (TLA 120.2 rotor) for 45 min at 4 °C. The supernatant was recovered and aliquots were flash frozen in liquid nitrogen and stored at −80 °C until use.

*Sp*Act1 and *Sp*Act1^H73me^ (Figures 1, 2, 3, 4, 5, S2, S4, S6, S7, S8, S9, S10)

*Schizosaccharomyces pombe* recombinant actin was expressed in and purified from yeast *Pichia pastoris* (*P. pastoris*) following published protocol (Hatano et al., 2018, 2020), yielding N-terminally acetylated actin *Sp*Act1 and both N-terminally acetylated and His73 methylated actin *Sp*Act1^H73me^, respectively. We note that *P. pastoris* has been reassigned to the *Komagataella* genus (Kurtzman, 2009).

Plasmid TH206 (pPICz-*Sp*Act1-Thymosinβ4-8xHis) was a kind gift from Pr. Balasubramanian. The Y371H mutation was introduced in the actin gene by site directed mutagenesis to avoid off target chymotrypsin cleavage (Tang et al., 2023), a mutation which has been shown to have no effect *in vivo* (Tang et al., 2023) using the KLD enzyme kit (New England Biolabs) and primers o393 (TGGTATCGTTCACCGTAAGTG) and o394 (GGTCCGCTCTCATCATAC) resulting in plasmid pIB0008 (pPICz-*Sp*Act1(Y371H)-Thymosinβ4-6xHis). This plasmid was linearized with PmeI then transformed by electroporation into *P. pastoris* strains MBY12334 (pIB2-NAA80 in *P. pastoris his4*) and MBY12805 (pIB4-myc-SETD3:Tcyc1-Pgap:NAA80:Taox1 in *P. pastoris his4*), kind gifts from Pr. Balasubramanian, yielding *Pp*GRAJ015 (pPICz-SpAct1(Y371H)-Thymosinβ4-8xHis pIB2-NAA80 in *P. pastoris his4* ; result *Sp*Act1) and *Pp*GRAJ023 (pPICz-SpAct1(Y371H)-Thymosinβ4-8xHis pIB4-myc-SETD3:Tcyc1-Pgap:NAA80:Taox1 in *P. pastoris his4* ; result *Sp*Act1^H73me^), respectively. Positive transformants were selected in YPD + 1000µg/mL Zeocine agar plates and validated by PCR with primers o175 (GACTGGTTCCAATTGACAAGC) and o176 (GCAAATGGCATTCTGACATCC) and stored in YPD + 25% glycerol at −80°C.

For actin production, *Pp*GRAJ015 and *Pp*GRAJ023 were inoculated into 1L MGY (1.34% YNB, 1% glycerol, 0.00004% biotin) and incubated O/N at 30°C 280rpm. Cells were then diluted in 6L MGY and incubated at 30°C 280rpm until OD600 reached 1.5. Media was then shifted to MM (1.34% YNB, 0.05% methanol, 0.00004% biotin) following centrifugation at 7500 rpm (F10-4×1000 LEX rotor) for 10min and washing with sterile water and cells were incubated 24h at 30°C, 280rpm. 0.5% methanol was then added to the cells, which were further incubated 24h at 30°C, 280rpm. Cells were then pelleted at 7500rpm for 10min, washed with water, and resuspended into 30mL sterile water. This slurry was used to freeze beads in liquid nitrogen, which were broken using a retsch with 20×1min cycles. The resulting powder was stored at −70°C until use, where it was resuspended in 2X Binding Buffer (20 mM Imidazole pH7.4, 20 mM Hepes pH7.4, 600 mM NaCl, 4 mM MgCl2, 2 mM ATP, 1 for 50mL EDTA-free Protease cocktail inhibitor tablets (Sigma), 1 mM PMSF, 7 mM β-mercaptoethanol). Lysate was sonicated then centrifuged at 4°C, 30000rpm (Ti45 rotor) for 1h and the supernatant was filtered through 0.45 µM membranes. The supernatant was then incubated under gentle agitation with 6mL equilibrated Ni-NTA agarose beads (Macherey-Nagel) for 2h. The resin was then poured into an econo-column (Bio-Rad), washed with 60mL 1X Binding Buffer and 90mL G Buffer (10 mM Tris pH7.8, 0.1 mM CaCl2, 0.01% NaN3, 0.2 mM ATP, 1 mM DTT). Chymotrypsin digestion was done O/N at 4°C in 20mL G Buffer at 10µg/mL. Digestion was stopped by addition of 2 mM PMSF for 30min at 4°C. The solution was allowed to flow through the econo-column and the beads were further washed with 15mL G Buffer. The elution containing the G-actin was collected, concentrated using a 30kDA cut off amicon and polymerized by addition of 10X KME (20 mM MgCl2, 50 mM EGTA, 1M KCl) 2h at room temperature then O/N at 4°C. The polymerized actin was pelleted by ultracentrifugation for 45min at 90000rpm (TLA 100.3 rotor), 4°C, and resuspended with a potter into 4mL G Buffer and depolymerized by minimum 3 rounds of dialysis in a 12-14kDa cut off servapor dialysis tube against 1L G Buffer for 24h, and further helped by sonication as needed. Dialyzed actin was then ultracentrifuged as before and the supernatant was collected and injected onto a gel filtration column. The pertinent fractions were identified as peak of fluorescence at 290nm and further validated by electrophoresis, pulled and concentrated using a 30kDA cut off amicon if needed. The left most fractions were excluded as they might contain actin dimers. Aliquots were flash frozen in liquid nitrogen and stored at −80°C until use.

*Sc*Act1 (Figures 1, 2, 3, 4, 5, S4, S5, S6, S10)

*Saccharomyces cerevisiae* recombinant actin was similarly expressed in and purified from *Pichia pastoris*. Plasmid TH3-37 (pPICz-ScAct1(Codon optimized for *P. pastoris*)-Thymosinβ4-8xHis) was a kind gift from Pr. Malasubramanian and was transformed into MBY12334 as before, yielding *Pp*GRAJ021. This strain was then used for production and subsequent purification as before, yielding N-terminally acetylated actin.

*Hs*ACTB (Figure S5)

*Homo sapiens* recombinant β actin was similarly expressed in and purified from *Pichia pastoris*. The strain MBY12870 (pPICZc-UbD-HsACTB_2:375-thymosin β-8His pIB4-myc-SETD3:Tcyc1-Pgap:NAA80:Taox1 in *P.pastoris his4*) was a kind gift from Pr. Malasubramanian and was used for production and subsequent purification as before, yielding actin both N-terminally acetylated and His73 methylated.

Native *Sc*Act1 (Figure S5)

*Saccharomyces cerevisiae* native actin was purified using a DNAseI affinity column as previously reported (Goode, 2002).

Briefly, commercially purchased baker’s yeast (Kastalia, LESAFFRE, France) was pelleted and rinsed once in cold H2O, then resuspended in 200mL cold H2O per liter of culture to obtain a cell slurry. This slurry was used to flash freeze beads in liquid nitrogen. The beads are subsequently reduced to powder using a Waring blender, and stored at −80°C until use.

DNase I was resuspended to 20mg/mL in Coupling buffer (100 mM Hepes pH7.2, 20 mM CaCl2, 1 mM PMSF) and coupled to a pre-equilibrated Affi-Gel 10 Resin (Bio-Rad) for 4h at 4°C under agitation. The coupled resin was then poured onto an econo-column (Bio-Rad) and washed sequentially with 0.1M Tris pH 7.5, G-buffer (10 mM Tris pH 7.5, 0.5 mM ATP, 0.2 mM DTT, 0.2 mM CaCl2), G-buffer supplemented with 0.2M ammonium chloride and G-buffer. The column was stored at 4°C until use.

The cell powder was thawed into G-buffer supplemented with protease inhibitors (Protease Inhibitor Cocktail Set IV, Calbiochem) and centrifuged at 160 000g for 30min at 4°C.The resulting supernatant was filtered through a cheese cloth and loaded onto the DNase I column. The column was then washed sequentially with G-buffer supplemented with 10% formamide, G-buffer supplemented with 0.2M ammonium chloride and G-buffer. The actin was eluted with G-buffer supplemented with 50% formamide and dialysed against modified G-buffer (10 mM Tris pH 7.5, 0.5 mM ATP, 0.2 mM DTT, 0.1 mM CaCl2) overnight. The actin was aliquoted and flash frozen in liquid nitrogen.

Spectrin actin seeds (Figures 1, 2, 3, 4, S2, S4, S5, S6, S9, S10)

Human spectrin actin seeds were extracted from red blood cells as previously published (Casella et al., 1986).

A pocket of packed red blood cells was obtained from Etablissement Français du Sang and dispatched in 50mL falcons pre-filled with 25mL Resuspension Buffer (5 mM NaPO4 pH7.7, 150 mM NaCl, 1 mM EDTA), then centrifuged at 5000g for 15min at 4°C. The resulting pellets were washed by two rounds of centrifugation in Resuspension Buffer, resuspended and pooled in 50mL Resuspension Buffer, then added to 700mL Lysis Buffer (5 mM NaPO4 pH7.7 + 1 mM PMSF), leading to the formation of ghost cells. The mixture was agitated for 40min at 4°C then centrifuged at 50000g for 15min at 4°C. The clear supernatant was carefully removed and the ghost cells resuspended by gentle manual rotation, which allows separation from the non-lysed red cell pellet. The resuspended ghosts were pulled together, diluted to 360mL in Washing Buffer (5 mM NaPO4 pH 7.7, 0.1 mM PMSF) and centrifuged at 50000g for 15min at 4°C. The previous operations were repeated twice with180mL Washing Buffer to wash the ghost cells. The final pellet was resuspended in 60mL Extraction Buffer (0.3 mM NaPO4 pH7.6, 0.1 mM PMSF) and centrifuged at 50000g for 30min at 4°C. The resulting pellet was resuspended in 60mL Extraction Buffer and incubated for 40min at 37°C with regular manual agitation, then ultracentrifuged at 400000g for 30min at 4°C. The supernatant was recovered and supplemented with 2 mM DTT, 1/200x proteases inhibitors (Sigma), then diluted in equal volume of cold Ethylene Glycol, and stored at −20°C until use. Concentration was assessed using a pyrene-actin assay to determine the concentration of growing barbed ends.

## Experiments

Microfluidics Experiments (Figures 1, 2, 3, 4, S2, S4, S5, S6, S9, S10)

All experiments were performed as previously published (Wioland et al., 2022) using a Poly DiMethyl Siloxane (PDMS, Sylgard) chamber, 20 µm in height, 1600 µm in width, 1cm in length, with 3 inlets and one outlet. Chambers were mounted on coverslips that were cleaned beforehand with a 4-step 30 minutes ultrasonic bath routine (2 mM Hellmanex at 35°C, 2 M KOH, milli-Q H2O, 100 % absolute ethanol). Chamber and coverslip were exposed to plasma for 40 s, bound together and heated at 100°C for 5 minutes. The chamber inlets were connected to tubes in which we control the pressure using a MFCS-EZ device (Fluigent), and the resulting flow rate was measured using Flow units (Fluigent). As a result, we can control and change very quickly the solution we expose the chamber to (Wioland et al., 2022).

Before all experiments, the chamber was connected in F-buffer (10 mM Tris pH 7.4, 1 mM MgCl2, 0.2 mM ATP, 0.2 mM EGTA, 10 mM DTT, 2 mM DABCO, 0.01 % NaN3, 50mM KCl), functionalized by injecting spectrin actin seeds (20 pM spectrin seeds in F-buffer for 3min), passivated by injecting 5% BSA for a minimum of 10 minutes and thoroughly rinsed out with F-buffer. Unless indicated otherwise, experiments were performed at 25 (±2) °C in F-buffer with 50 mM KCl. For Phosphate Buffer, Tris and KCl were replaced by 80.2% K2HPO4, 19.8% KH2PO4 (proportions leading to a pH7.4 solution), total concentration as indicated. For Sulfate Buffer, Tris and KCL were replaced by the indicated concentration of Na2SO4, and the pH adjusted with HCl. Unless indicated otherwise, actin was labelled with a half molar ratio of N6-(6-Aminohexyl)-ATP-ATTO-488 (Jena Bioscience). In that case, F-buffer without ATP was used. Because the ATP concentration in the actin Storage Buffer is always 200 µM but the actin concentrations depend on the specific actin prep, the obtained labelling ratio when using a half-molar ratio of ATP-atto488 relative to actin differ but were always between 4.3 and 7.7%. Concentrations of proteins flowed in are specified in the figures

Fixed filament experiments (Figure 5)

For experiments pertaining to persistence length measurements, 6 µM of the appropriate Ca-ATP-G-actin originally in G-buffer (2 (OcACTA1) or 10 mM (SpAct1, ScAct1, SpAct1^H73me^) Tris pH7.8, 0.1 mM CaCl2, 0.01% NaN3, 0.2 mM ATP, 1 mM DTT) were first incubated in ME buffer (15 µM MgCl2, 0.2 mM EGTA) and the appropriate concentration of ATP-atto488 so that we reach a 50% labelling ratio of the actin monomers (i.e. a concentration equal to the one of ATP, which depends of how much we diluted the actin) for 10min at RT. The aim is to obtain actin where the cation and ATP-atto488 exchange has had enough time to proceed so that we obtain fluorescently homogenous filaments. Next, the Mg2+-ATP-G-actin was supplemented with 50 mM KCl, and polymerized at 5 µM for 1h at RT, after which we can assume all of the F-actin is in the ADP state.

The Mg-ADP-F-actin were diluted to 0.0005 µM with in F-buffer without ATP (10 mM Tris pH 7.4, 1 mM MgCl2, 0.2 mM EGTA, 10 mM DTT, 2 mM DABCO, 0.01 % NaN3, 50 mM KCl) and supplemented with 0.1 µM of the appropriate unlabelled Ca2+-G-actin to prevent depolymerization. A 20µL drop of the dilution was deposited onto a coverslip which had been plunged into a 0.002mg/mL poly-l-lysine solution for 2s and dried with compressed air. This concentration of poly-l-lysine allows for a slow deposition of the filaments so that we can extract a persistence length as close as possible of freely fluctuating filaments. After waiting 2/3min for the settling of the filaments, multiple snapshots were taken inside of the droplet away from the border as quickly as possible to avoid drying of the droplet.

Open Chamber experiments (Figure S7)

22×40 and 18×18 mm coverslips were first passivated using 200 or 70µL, respectively, of a 1mg/mL Peg-silane solution in 95% Ethanol and 0.1% HCl which was dropped on the coverslip, then dried at 70°C for 18min. The coverslips were rinsed sequentially with 100% Ethanol, 70% ethanol and mqH2O. The chamber was mounted using parafilm strips and the two coverslips oriented perpendicularly, forming 4 canals (Figure S7A), and heated while being pressed down on a hot plate for a short time so that the canals become impervious. After mounting the chamber on the microscope, solutions are injected on one side of the chamber and flow is induced by aspiration with blotting paper on the other side. Canals are sealed using Vitrex and acquisition is started immediately.

To follow actin polymerization, canals are first injected with 100µL of F-buffer without ATP supplemented with 0.18% of methylcellulose 4000 cp, then with 100µL of 0.35µM of the appropriate actin labelled with a half molar ratio of ATP-atto488 in the same buffer. A movie with a 10s interval is acquired.

### Image Acquisition and Analysis

Image acquisition

The microfluidic chamber (or the open chamber or the coverslip) was attached on a motorized stage (Marzhauser) onto a Nikon Ti2 inverted microscope, equipped with a TIRF 1.49 NA 100 x oil-immersion objective, a sCMOS-kinetix camera (photometrics), a dichroic Quad band and emission filter ZET405/488/561/647 cube, tunable 488, 561 and 642 nm 100 mW lasers, which were controlled using a Total Internal Reflection Fluorescence (TIRF) set-up (iLAS2, Gataca Systems) through the micromanager software (Edelstein et al., 2010; D. Edelstein et al., 2014). Please note that depending on the passivation quality, the background fluorescence can vary. As a result, illumination power and exposure time could vary between experiments, but were usually set around 10% and 150ms, respectively. The time interval between frames was adapted depending on the experiment.

Linear polymerization/depolymerization rates measurements (Figures 1E, 1F, 1G, 1H, 1I, 2C, 2D, 4B, 4C, S4, S5, S6)

For each experiment, a line was traced over 30 randomly selected filaments and a kymograph was obtained for each of them and rotated such that time corresponds to the abscissa and the length of the filament the ordinate, pointed end at the bottom. The angle θ in degree formed by the line following the filament barbed end over time (yellow line in Figure 1D) and the abscissa was recorded for each of those kymographs, and the corresponding polymerization/depolymerization rate V was calculated based on the following formula:

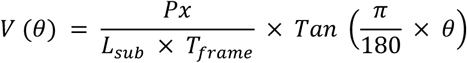

With Px being the pixel size in nm, L_sub_ being the length added to a filament by the addition of one actin subunit in nm, fixed to 2.7nm, and T_frame_ being the time interval between the different frames. The 30 obtained values were averaged to obtain one rate per replicate, which depending on the experiment were either plotted as mean of all replicates and standard deviation between replicates (Figures 1E, 2C, 4B-C) or as individual replicate averages with a black bar displaying the mean of all replicates when present (Figures 1F, 1G, 1H, 1I, 2D, S4, S5, S6). For depolymerization experiments, the first few frames were excluded from the analysis so as to bypass the intermediary ADP·Pi state. Filaments which are blocked due to non-specific interaction with the surface (detectable by the absence of the small transverse oscillations observable at low flow) were excluded from the analysis.

Affinity of Pi for the barbed end (Figures 2C and 4C)

The barbed end depolymerization of filaments is measured in the presence of Pi in the buffer. Considering that Pi rapidly binds and unbinds the nucleotide pocket at the barbed end, we can write the depolymerization rate as v_depol_ = k_off_^ADP^ + (k_off_^ADP·Pi^-k_off_^ADP^) x [Pi]/(K_D_^Pi, BE^ + [Pi]) , where K_D_^Pi, BE^ is the equilibrium dissociation constant of Pi at the barbed end, and where k_off_^ADP·Pi^ and k_off_^ADP^ are the off-rate constants of ADP·Pi- and ADP-actin, respectively.

In addition to binding the nucleotide pocket of actin, Pi may affect the barbed end depolymerization rate through ionic strength and Hofmeister effects. At high [Pi], this seems to cause an additional decrease of the depolymerization rate (right panels in Figure S4). To mitigate this effect, we chose to set k_off_^ADP·Pi^ to the values we measured at 200 mM Pi (yeast actins) and 150 mM Pi (rabbit actin). The equilibrium dissociation constant K_D_^Pi, BE^ and the off-rate constant k_off_^ADP^ were the free parameters when fitting the depolymerization rate vs. [Pi] with the above hyperbolic equation, in Figures 2C and 4C, using least-square minimization (Scipy python package). The resulting values for k_off_^ADP^ matched our independent measurements (Figures 1E, F and 4B) and the values measured in the absence of Pi in Figures 2C and 4C.

Pi-release rate measurements (Figures 3D-G, 4D, S9)

For those experiments, actin was first polymerized in phosphate buffer, leading to filaments fully in the ADP·Pi state (Figure 3A). Filaments were then exposed to a buffer without phosphate nor actin, so that they start both to depolymerize and to release their phosphate. The depolymerization rate is linked to the phosphate content, so that the depolymerization rate is not constant over time (Figure 3B) and we measured the instantaneous depolymerisation rate over time for such experiments (Figure S9). This was done using a custom Python script (Kotila et al., 2022; Schahl et al., 2025). This algorithm tracks the barbed end of each filament over time and extracts the instantaneous depolymerization rate at time *t* by using a linear fit over 5 frames centered on time *t*. The data of at least 40 filaments was pooled and the instantaneous depolymerization rate v_depol_(t) is then fitted as a function of time to the following equation using curve-fit Scipy function:

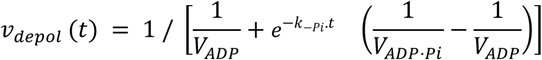

where V_ADP_ is the depolymerization rate of ADP-actin filaments, V_ADP·Pi_ the depolymerization rate of ADP·Pi-actin filaments and k_-Pi_ the rate of Pi release, which are free parameters. The values of the parameters obtained for each replicate are plotted as individual values (circles) and mean (black bar).

Because at the end of the experiment, the filament is fully in the ADP-state, V_ADP_ is in agreement with k_off_^ADP^ (see Figure 4D vs Figures 1F, 4C and Table 3). However, because the Pi release is faster at the barbed end than the rest of the filament, there are two routes for an ADP·Pi-actin subunit to dissociate from the barbed end (Figure 4C): either the ADP·Pi-actin subunit dissociates as is, with the k_off_^ADP·Pi^ rate, or it first releases its phosphate with a k_-Pi_^BE^ rate and then dissociates with a k_off_^ADP^ rate, so that V_ADP·Pi_ does not match k_off_^ADP·Pi^, but instead can be used to extract k_-Pi_^BE^ (Table 3) with the following equation:

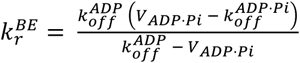

Persistence length measurements (Figure 5C-D)

Stacks of snapshots of poly-l-lysine adhered ATP-atto488 labelled filaments were background subtracted on FIJI (Schindelin et al., 2012) with a rolling ball radius of 20 pixels, skeletonized using TSOAX (T. Xu et al., 2019) with default parameters except for Ridge Threshold (0.035), Minimum Foreground (60), Minimum SOAC Length (25), Maximum Iterations (1000), Alpha (0.05) and Beta (0.16). The resulting snakes were then processed using a custom made Python algorithm. The tangent angle along the skeletonized filament was measured every 520 nm (every 4 pixels) and the angular cosine correlation was quantified between two points along the filament, as a function of their distance δs. For each actin species, data were collected from at least 11 independent repeats, with a minimum of 323 filaments (average length from 4.6 to 7.95 µm). Data were then fitted by the theoretical 2-dimensional angular correlation of a filament of persistence length Lp:

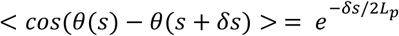

using the ‘curve-fit’ from the Scipy python package and are shown in Figure 5C. Shaded areas are standard errors of the fit. Persistence Lengths obtained for max curvilinear distances 5.72 and 6.24 were calculated and averaged for each replicate and are shown in Figure 5D.

Intensity along the filament in open chamber (Figure S7D)

Roughly 10 minutes after polymerization started, a broken line was traced along the length of 20 filaments, going beyond the edges, and the intensity of the filament is recorded for each pixel along this length. Each intensity profile is normalized by its maximum and oriented by its brightest end. For *Oc*ACTA1, this will correspond to the barbed end. For *Sp*Act1, this is essentially random. The beginning of the filament is identified as the pixel where it reaches half maximum intensity and profiles are truncated and aligned at this end. Profiles are truncated at the other end at 7µm, a length which is shorter than the shortest filament so that we are still inside of the filament along the whole profile. Intensity profiles were averaged and plotted in Figure S7D with error bars being standard deviations.

Plot rendering and statistical analysis

Except for Figure 5C, S8 and S9, all plots and statistical analysis were made using Graphpad Prism v10.1.2.

Symbols in Figure 1F, 2D, 3D-G, 4D and 5D refer to the result of an unpaired student t-test (ns : P>0.05 ; * : P≤0.05 ; ** : P≤0.01 ; *** : P≤0.001 ; **** : P≤0.0001)

### Phylogenetic analysis

Uniprot sequences for *Sc*Act1 (P60010), *Sp*Act1 (P10989), *Oc*ACTA1 (P68135) and *At*ACT1 (P0CJ46), which as taken as an external group, were aligned and a phylogenetic tree was obtained using ngphylogeny.fr (Lemoine et al., 2019; Junier & Zdobnov, 2010; Katoh & Standley, 2013; Criscuolo & Gribaldo, 2010; Guindon et al., 2010; Lemoine et al., 2018)

The tree display was visualized and manipulated to our convenience with SeaView (Gouy et al., 2010). Unit is the average number of substitutions per site.

The alignment displayed in Figure S1 was obtained using the uniprot sequences listed in the legend using clustalOmega (Madeira et al., 2024). It was then formatted for publication using JalView (Clamp et al., 2004) using clustal color coding and 30% conservation cut-off.

### Actin Structure Rendering

Structural sketch of ADP-F-actin from *S. Cerevisiae* shown in figure 4A is adapted from PDB 9GO5 (Stevenson et al., 2025), using ChimeraX (version 1.9 (2024-12-11)).

### LC-MS of SpAct1^H73me^ (Figure S8)

Samples (200 ng) were injected onto a custom made BioResolve RP (450Å, 2.7 µm, 0.3 mm x 150 mm) Polyphenyl Column (Waters, Milford, MA, USA) using an Acquity M-Class chromatographic system from Waters. Mobile phases A and B were H_2_O and H_2_O/ACN 20/80, respectively, acidified with 0.1% (v/v) DFA. Samples were eluted for 35 min using the percentage of acetonitrile described in the table below. The eluent was sprayed using the conventional ESI ion source of a Waters Cyclic IMS SELECT SERIES mass spectrometer operating in the positive ion mode. The quadrupole was operated in RF-only mode, the TOF was operated in V-mode with a resolution specification of 60,000 FWHM at a mass range of m/z 50-4000. Data were deconvoluted using the Intact Mass v5.3.44-gbc4a1db565 software provided by Protein Metrics.

| Time (min) | Flow (µL/min) | %B |
| --- | --- | --- |
| 0 | 10 | 10 |
| 3 | 10 | 10 |
| 4 | 6 | 13.9 |
| 5 | 6 | 17.6 |
| 6 | 6 | 20.4 |
| 7 | 6 | 22.7 |
| 8 | 6 | 24.7 |
| 9 | 6 | 26.4 |
| 10 | 6 | 27.9 |
| 11 | 6 | 39.2 |
| 12 | 6 | 30.4 |
| 13 | 6 | 31.5 |
| 14 | 6 | 32.5 |
| 15 | 6 | 33.8 |
| 16 | 6 | 34.8 |
| 18 | 6 | 36.7 |
| 20 | 6 | 38.6 |
| 22 | 6 | 40.7 |
| 24 | 6 | 43.3 |
| 26 | 6 | 46.6 |
| 28 | 6 | 51.3 |
| 29 | 10 | 60 |
| 31 | 10 | 60 |
| 32 | 10 | 10 |
| 35 | 10 | 10 |

## Notes

### Competing Interest Statement

The authors have declared no competing interest.

### Summary of Updates

Compared to v2, new data and analysis have been added, some points have been clarified.

